# Cardiac Cycle Modulates Alpha and Beta Suppression during Motor Imagery

**DOI:** 10.1101/2024.04.03.587974

**Authors:** Giuseppe Lai, David Landi, Carmen Vidaurre, Joydeep Bhattacharya, Maria Herrojo Ruiz

## Abstract

**Introduction:** The baroreceptor hypothesis posits that baroreceptors, located on the cardiac walls, are most active during systole, translating cardiac contraction information to the brain. Studies within this context have suggested that the systolic phase, characterised by increased noise, may compromise the processing of sensory stimuli. Although the effect of systolic and diastolic cardiac cycle phases on cognition, perception, and action has been widely documented, there remains a gap in applying these interoceptive insights to enhance assistive technologies such as brain-computer interfaces (BCIs). In the context of BCIs, motor imagery (MI) —the mental rehearsal of movement—serves as a widely used control paradigm, yet its modulation through the cardiac cycle has not been empirically tested. Bridging this gap, this study examined how the cardiac cycle phases influence MI by assessing their effect on contralateral suppression of alpha (8-13 Hz) and beta (14-30 Hz) activity in primary sensorimotor cortices.

**Materials & Methods:** Twenty-nine participants performed left/right thumb abductions based on the direction of an arrow presented on the screen to get familiarised with kinesthetic sensations. They then completed a MI task of the same movements. We recorded both electroencephalography (EEG) and electrocardiography (ECG), focusing our analysis on data epochs aligned with the experimental cue, based on whether it occurred during the systolic or diastolic phase of the cardiac cycle. Time-frequency analysis of source-reconstructed data assessed cue-induced changes in power spectral density (PSD) within the alpha and beta bands in the postcentral and precentral gyrus.

**Results:** We found that alpha and beta suppression in the contralateral primary motor and somatosensory cortex was more pronounced when the cue fell during the diastolic phase of the cardiac cycle than during the systolic phase. Validating the main results, an analysis with circular statistics revealed that trials with particularly pronounced contralateral alpha and beta suppression featured cues with latencies clustering during diastole, the quietest time of the cardiac cycle. Accompanying the EEG effects, EMG activity on the side of the movement was enhanced during diastole.

**Conclusion:** These findings provide evidence that MI performance can be enhanced by considering the cardiac cycle phases, offering promising implications for BCI-based applications.

- ***Key point 1:*** *The phases of the cardiac cycle influence motor imagery performance*.
- ***Key point 2:*** *Alpha and beta contralateral suppression over sensorimotor cortices is more pronounced when movement direction is cued during diastole*.
- ***Key point 3:*** *Contralateral suppression is more likely to cluster during the quietest time of the diastolic phase: between the T-wave and the P-wave*.

## Introduction

Interoception refers to the ability of the central nervous system (CNS) to sense, interpret, integrate, and regulate information emerging from visceral organs, such as the heart, gastrointestinal tract, and respiratory system (Barrett & Simmons, 2015; Craig, 2009; Critchley & Garfinkel, 2018; Park et al., 2020). This interplay has been shown to significantly influence how we perceive the external world, shaping our cognitive, perceptual, and emotional experiences (Azzalini, Rebollo, & Tallon-Baudry, 2019). The relationship between internal organs and the CNS is bidirectional, involving descending and ascending pathways to regulate visceral functions and transmit organ-related information to higher-order cognitive areas (Azzalini et al., 2019). In the case of the heart, this phenomenon is known as heart-to-brain interaction (HBI) (Engelen, Solcà, & Tallon-Baudry, 2023).

At each cardiac cycle, blood fills the atria during ventricular diastole, when the heart is most relaxed, and is then ejected into the circulatory system during ventricular systole, when the ventricles are most contracted. It has been suggested that baroreceptors found on cardiac walls (including the atria, ventricles, and aortic arch) inform the brain about the timing and strength of cardiac contractions (Azzalini et al., 2019; Duschek, Werner, Reyes del Paso, 2013; Engelen et al., 2023). This information travels via the vagus nerve and reaches subcortical brain structures (such as the brainstem, and thalamus) where it is processed before being relayed to central cortical areas, including the prefrontal cortex, anterior and posterior insular cortex, cingulate cortex, and primary and secondary somatosensory cortex (Azzalini et al., 2019; Engelen et al., 2023). The resulting influence of this ascending process can be measured by synchronising neural and cardiac activity recorded via electroencephalogram (EEG) and electrocardiogram (ECG), respectively. A well-known approach involves finding the relationship between the timing of experimental cues and the cardiac cycle, analysing neural responses separately for systole and diastole (Bury, García-Huescar, Bhattacharya & Ruiz, 2019; Grund et al., 2022; Park & Tallon-Baudry, 2014; Park, Correia, Ducorps, Tallon-Baudry, 2014; Pramme, Larra, Schächinger, & Frings, 2014; Pramme, Larra, Schächinger, & Frings, 2016).

Research on HBI has demonstrated that the cardiac cycle modulates perception (Barrett & Simons, 2015; Schulz et al., 2009), cognition (Garfinkel et al., 2014; Honda & Nakao, 2022), affective processing (Critchley & Garfinkel, 2017; Gray et al., 2012), and action (Al et al., 2023). This evidence has been particularly relevant for psychiatric research, where mental health conditions are associated with imbalances in brain-body interaction (Nord & Garfinkel, 2022; Khalsa et al., 2018; Khoury, Lutz, & Schuman-Olivier, 2018). However, there is a lack of translational research that investigated these insights for applications in other clinical settings, such as neurological conditions, and in assistive technologies for neuromuscular disorders. Motor imagery (MI)—rehearsing and focusing on the kinesthetic sensations of a movement (Savaki & Raos, 2019)—stands as one of the main paradigms traditionally used in brain-computer interface (BCI) studies (Sannelli, Vidaurre, Muller, & Blankertz, 2019), as well as a common rehabilitative exercise for post-stroke survivors (Khan, Das, Iversen, & Puthusserypady, 2020; Tong et al., 2017). Yet, the interplay between interoception and sensorimotor processes lacks sufficient evidence, particularly regarding MI. Understanding the interaction between the cardiac cycle and sensorimotor cortices during imaginary movements may uncover optimal time windows for MI performance modulation, leading to innovative approaches for improving MI-based assistive technologies.

This study investigates the modulation of neural sensorimotor activity during MI as a function of the cardiac cycle. Two prevailing hypotheses propose contrasting effects during the most active phase of the cardiac cycle, depending on the type of processing involved—sensory or motor. One suggests facilitation, while the other indicates detrimental effects. The baroreceptor hypothesis (Lacey, 1967) posits that during systole, when baroreceptors are most active, the signal-to-noise ratio in the CNS is significantly reduced, leading to decreased cortical excitability and attenuated sensory processing (Duschek et al., 2013; Rau, Pauli, Brody, Elbert, & Birbaumer, 1993). Recent studies align with the baroreceptor hypothesis, showing reduced cortical excitability and diminished sensory processing across various modalities. For instance, aversive auditory stimuli at systole elicited weaker startle responses (Schulz et al., 2009). Moreover, stimuli in visual, auditory, and tactile domains presented during systole resulted in slower reaction times (Edwards et al., 2007) and required extended intervals to sample sensory information during self-initiated touches (Galvez-Pol et al., 2022). Systolic electrical stimulation in the somatosensory realm reduced pain thresholds and stimulus perception (Grund et al., 2022; Motyka et al., 2019). However, findings in the visual domain are mixed: while some research indicated enhanced visual discrimination and search at systole (Pramme et al., 2014; Pramme et al., 2016), another study reported no impact on visual discrimination during systole (Park & Tallon-Baudry, 2014; Park et al., 2014).

In the motor domain, a complementary hypothesis, supported by further experiments, proposes a facilitatory effect on actions initiated at systole. For instance, Rae et al. (2018) found that response inhibition in a stop signal task was more efficient during systole, evidenced by shorter reaction times. Systolic phases, marked by higher cardiovascular arousal, have been linked to increased self-paced oculomotor activities, such as microsaccades (Ohl et al., 2016). Additionally, Kunzendorf et al. (2019) observed that participants were more likely to initiate the presentation of visual stimuli during systole, implying that active environmental sampling is favoured at this time.

Furthermore, recent research demonstrated that involuntary motor-evoked potentials (MEPs), triggered by transcranial magnetic stimulation (TMS), were larger during systole, indicating motor facilitation amidst higher cardiovascular arousal (Al et al., 2023). This study also observed that desynchronisation in the alpha and beta frequency bands (8–25 Hz) was significantly more pronounced at systole than diastole. However, some earlier studies failed to replicate these findings (Bianchini et al., 2021; Filippi et al., 2000; Otsuru et al., 2020). Moreover, additional findings in the motor domain revealed mixed effects regarding the facilitatory role of systole on action. For example, non-elite rifle shooters often fire during systole (Konttinen et al., 2003), whereas elite shooters show a preference for diastole (Helin et al., 1987). In addition, a recent application of real-time ECG analysis for assessing response execution during cognitive-emotional control demonstrated faster responses when stimuli were presented at diastole and associated with enhanced theta-band (4–7 Hz) activity (Adelhöfer, Schreiter, & Beste, 2020).

Given the potential implications in the field of EEG-based BCIs, understanding the role of cardiac interoception in MI is crucial, and this includes determining its alignment with either the baroreceptor hypothesis or the proposal of motor facilitation during systole. MI in laboratory settings typically involves the processing of cues and their internal rehearsal, a process that could be influenced by the effects of the cardiac cycle during both cue processing and the onset of imagined movements.

Both real movement execution (motor execution, ME) and its imagination activate the primary somatosensory cortex (S1) and the primary motor cortex (M1) (Grèzes & Decety, 2001; Hardwick, Caspers, Eickhoff, & Swinnen, 2018; Hetu et al., 2013; Lotze et al., 1999), with studies reporting overlapping activation in M1. Given that interoceptive information is integrated at central cortical levels, affecting both somatosensory and motor processes (Azzalini et al., 2019; Al et al., 2023), our primary hypothesis is that neural responses during MI—indexed by contralateral alpha and beta modulation in M1 and S1—are modulated by the phase of the cardiac cycle. Moreover, we anticipated that MI initiated following cues in the quiet phase of the cardiac cycle, diastole, would result in enhanced performance, reflected by more pronounced contralateral alpha and beta suppression, aligning with the baroreceptor hypothesis.

To test these hypotheses, we conducted an experiment where 29 participants performed a ME task, followed by a MI task involving the same movements, while we recorded their synchronised EEG and ECG activity. These tasks required participants to perform or imagine left and right thumb abductions, guided by the direction of an arrow displayed on the screen. Our results reveal that imaginary movements instructed during diastole lead to more pronounced alpha and beta suppression in contralateral sensorimotor cortices. This is accompanied by increased EMG activity on the side of the imagined movement.

## Materials & Methods

### Participants

The study was conducted at Goldsmiths, University of London, and received ethical approval from the local ethics committee at the Department of Psychology. The sample comprised 29 right-handed healthy volunteers (16 females), aged between 18 and 40 years (mean age = 20.20 years, standard error of the mean, SEM = 0.66). Participants received cash incentives or course credits for taking part in the study. Eligibility criteria included right-handedness, normal or corrected-to-normal vision, and no known history of neurological conditions. We aimed for a sample of 30 participants but excluded one post-experiment upon their disclosure of a neurological disorder, which disqualified them from the participation criteria. This sample size was informed by published effect sizes for cardiac influences on motor performance (Azevedo, Garfinkel, Critchley, & Tsakiris, 2017; Bury et al., 2019), which suggested Cohen’s *d* in the range 0.55–0.7 (probability of superiority of 0.69, equivalent to Cohen’s *d* ∼ 0.7 in Bury et al 2019). This implies that 24–28 participants would be sufficient to detect within-subject behavioural effects with 0.8 power.

### Experimental design

Participants were seated in a dimly lit room, facing a computer screen which displayed instructions and experimental stimuli. The experimental paradigm consisted of two tasks, preceded by an eyes-open resting-state recording lasting 5 minutes and an auditory oddball task (not analysed here). The experimental paradigm was programmed using PsychoPy (Peirce et al., 2019), a Python-based open-source software toolbox designed for the presentation of visual and auditory stimuli (Python v. 3.7.11; PsychoPy v. 2021.1.4; Ubuntu v. 20.04.5 LTS - Focal Fossa).

### Motor Execution (ME) and Motor Imagery (MI)

The motor execution (ME) and motor imagery (MI) tasks were modelled on the protocol established by Sannelli et al. (2019). In the ME task, participants executed a thumb abduction, either left or right, in response to the direction of an arrow displayed on the computer screen after a fixation cross. The abduction required participants to gently lift their thumb for approximately one second, maintaining it at maximum extension until the arrow reverted to the fixation cross, which occurred four seconds after the arrow. Importantly, participants were asked to focus their attention on the kinesthetic aspects of the movement such as muscle contraction and movement velocity. The kinesthetic sensation was defined as any sensation experienced during the execution of the movement that the participant felt salient. In the MI task, participants were instructed to mentally recreate the kinesthetic experience of the movement without physical execution or visualisation, ensuring consistency in sensations and similar timing throughout the task.

As shown in **Figure 1a**, each trial lasted approximately 8 seconds. The ME task consisted of a single block of 50 trials (25 left and 25 right), with an automatic 15-second pause after the first 25 trials. For the MI task, participants completed two blocks of 100 trials each, with an automatic 15-second pause every 25 trials and a break between the two blocks. The number of left and right trials was evenly distributed (100 trials for each condition) and their sequence was randomised.

**Figure 1.**
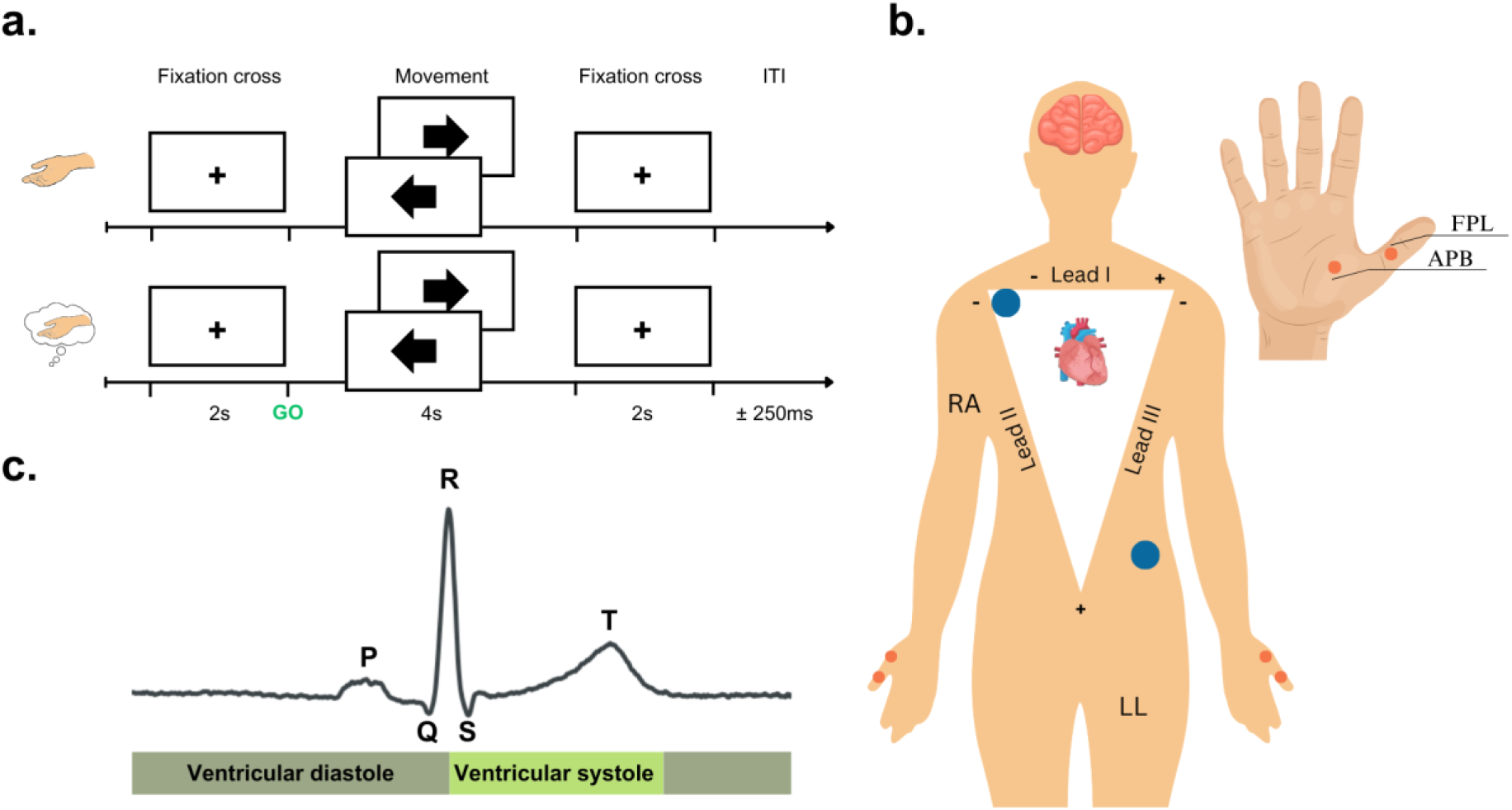
Illustration of the experimental paradigm. **a.** The experiment consisted of the motor execution (ME) task, where participants performed either a left or right thumb abduction movement based on the direction of the arrow and that lasted for approximately one second after the presentation of the cue/arrow. For the subsequent three seconds, participants were required to maintain the thumb in the lifted position and then relax once the fixation cross reappeared on the screen. **b.** The diagram shows the locations of the ECG electrodes (indicated by blue dots) based on the lead-II system configuration, and the placements of the EMG electrodes (shown as orange dots). One electrode was placed on the flexor pollicis longus (FPL) and the other one on the abductor pollicis brevis (APB). **c.** A schematic representation demonstrating the key events in the ECG trace, along with definitions for the phases of the cardiac cycle, specifically the ventricular diastole and ventricular systole.

### EEG, ECG, EMG recordings

EEG data were collected using the BioSemi ActiveTwo (BioSemi Inc.) with a 64-channel layout based on the 10-20 electrode-placement system and were recorded at a sampling rate of 1024 Hz. Two external electrodes were positioned on the left and right mastoids as an initial reference upon importing the data. ECG data were recorded using the lead II configuration: the negative electrode was positioned under the right collar bone, and the positive electrode was placed above the left hip bone (**Figure 1b**). Additionally, EMG data from both the left and right thumbs were recorded using 4 bipolar electrodes, positioned on the respective abductor pollicis brevis muscle of each thumb (**Figure 1b)**. Due to low signal-to-noise ratio in the EMG signal of three participants, their data was excluded from the main analysis. All signals were recorded using a high-pass filter at 0 Hz and a low-pass filter at 208 Hz.

### EEG data preprocessing

Preprocessing, epoching and artefact rejection were carried out using the open-source Python library: MNE-Python (Gramfort et al., 2013) (Python v. 3.9.7, MNE-Python v. 1.3.0, Ubuntu v. 20.04.5 LTS - Focal Fossa). The EEG data were re-referenced to common-average reference and down-sampled at 256 Hz. They were then notch-filtered (zero-phase, FIR design) at 50 Hz and its harmonics, and band-pass filtered (zero-phase, FIR design) between 0.5 Hz and 40 Hz. For both ME and MI trials, data were time-locked to the onset of the arrow (or cue) on the screen and epoched between -1 and 4 seconds relative to the cue onset. The data were first visually inspected for rejecting large artefacts, and bad channels were interpolated using spherical splines (Perrin, Pernier, Bertrand, & Echallier, 1989). Finally, independent component analysis, ICA (fastICA, Hyvärinen & Oja, 2000) was used to remove artefact components related to eye blinks, eye saccades, and cardiac artefacts. To ensure the removal of cardiac artefacts from EEG signals in the case that no cardiac IC were identified, we used a regression-based method (https://github.com/Giuseppe-1993/HBI-motor_imagery). A related approach has also been recently proposed (Arnau, Sharifian, Wascher, & Larra, 2023).

### ECG data preprocessing

To remove various types of artefacts in the ECG data, such as power-line noise, muscle artefacts, electrode contact noise and low-frequency baseline drifts, filters were applied to the continuous ECG recordings. Specifically, a band-pass filter (0.2-40 Hz; zero-phase, FIR design) was employed to eliminate slow drifts and reduce the impact of muscle artefacts. To detect key cardiac events in the ECG data, we utilised NeuroKit2 (v. 0.2.4), an open-source Python library (Makowski et al., 2021). This enabled the identification of the P-wave, R-peak, and T-wave within the ECG recordings. After Azzalini et al. (2019), we defined the systolic phase as the ventricular systole, which extends from the R-peak to the T-wave offset, and the diastolic phase was defined as the ventricular diastole, spanning the interval from the T-wave offset to the next R-peak (**Figure 1c**).

### EMG data preprocessing

Following the recommendations set by Konrad (2005), EMG data were bandpass filtered between 10 and 500 Hz. To ensure a robust estimation of standard quantitative properties of the signal’s amplitude, such as mean, peak, minimum, and maximum, we applied a full wave rectification of the signal, which involved converting all negative values to positive ones. Additionally, we applied a smoothing function (root mean square, RMS) to retain only the mean power of the signal. This approach provided a reliable measure of muscle activation, both ipsilateral and contralateral, over time during the tasks of overt thumb abduction (ME task) and covert thumb abduction (MI task).

### ECG and EMG Data Analysis

Initially, we conducted an analysis of the ECG signal for sanity checks, examining if the average ECG profile varied between epochs associated with left-cue and right-cue, separately for ME and MI tasks. Additionally, we contrasted the inter-beat intervals (IBIs) of participants across the ME and MI tasks. In the analysis of EMG data, our primary focus was on identifying differences between ipsilateral and contralateral activations separately in each motor task. We hypothesised that (i) there would be a pronounced ipsilateral activation compared to contralateral activation during ME, and (ii) there would be no significant difference in activation during the MI task. Finally, based on a previous study that reported larger muscle activity at systole (Al et al., 2023), we assessed the differences between the ipsilateral EMG traces for trials initiated during systole or during diastole. This analysis was carried out separately for the ME and MI tasks.

### Source Reconstruction

Neural activity at the source level was reconstructed using the Linearly Constrained Minimum Variance (LCMV) beamforming technique (Sekihara & Nagarajan, 2008; Van Veen, Van Drongelen, Yuchtman, & Suzuki, 1997; Van Vliet, Liljeström, Aro, Salmelin, & Kujala, 2018). LCMV beamforming requires two main ingredients: the forward solution (involving head geometry and conductivity properties) and the covariance matrices (for data and noise). For the forward solution, we used a three-layer head geometry and conductivity model for a standard brain provided by MNE-Python, which employs the boundary element model method. The data-covariance and the noise-covariance matrices were estimated for each participant, utilising a specific time window of interest (500–2000 ms post-cue) for the former and a pre-cue time window (-1000 to 0 ms) for the latter. The window of interest was selected based on the instructions provided to participants, who were directed to initiate the movement within the first second following the presentation of the cue. (see **Figure 3** for the EMG activity). This procedure resulted in 61452 dipoles bilaterally. In an exploratory analysis, we expanded this analysis to the full window length of 4 seconds.

We chose to parcellate the brain using the Desikan–Killiany–Tourville (DKT) cortical atlas (Klein & Tourville, 2012), which is integrated into MNE-Python and comprises 68 brain regions bilaterally (34 per hemisphere). The motor areas in this atlas include the precentral gyrus (primary motor cortex, M1), postcentral gyrus (primary somatosensory cortex, S1), and paracentral gyrus. Based on previous research on motor imagery (Hardwick et al., 2018; Hetu et al., 2013), our regions of interest (ROI) from the DKT atlas were: the precentral gyrus (M1) and the postcentral gyrus (S1). To reduce dimensionality, the dipole-related time series were mapped to these ROI using the principal component analysis (PCA)-flip method of MNE-python, resulting in a single signal for each region and hemisphere.

The paracentral gyrus was excluded because it mainly represents lower limb movements as a combined extension of the M1 and S1 regions. The supplementary motor area (SMA) is anatomically located on the medial aspect of the superior frontal gyrus (SFG), and therefore there could be some overlap between the labelled SFG in the DKT atlas and the region where the SMA is functionally located. However, we did not include the SFG in our analysis because the SMA is not distinctly isolated in this atlas. The Destrieux Atlas (Destrieux, Fischl, Dale, & Halgren, 2010) includes the SMA, but is more suited for MRI/fMRI studies or EEG/MEG studies that utilise individual standard MRI images, due to its detailed labelling with 78 labels per hemisphere. This fine-grained labelling demands higher spatial resolution for accurate source reconstruction.

### Time-frequency decomposition

For the time-frequency analysis of EEG signals during MI and ME tasks, we employed a wavelet transform with Morlet wavelet in MNE-Python (Cohen, 2019; Tallon-Baudry, Bertrand, Delpuech, & Jacques Pernier, 1997). This analysis decomposed the EEG epochs into their constituent frequency components ranging from 8 and 30 Hz, specifically alpha (8–13 Hz) and beta (14–30 Hz) bands. The wavelet decomposition was performed on the entire duration of the epochs, spanning from -1 second to 4 seconds relative to the cue, using a setting of 5 cycles and a resolution of one bin per frequency (1 Hz). The resulting time-frequency power spectral density was baseline-corrected to the pre-cue interval using the z-score method in MNE-Python, between -0.5 and 0 seconds pre-cue, as the baseline interval. Therefore, our time-frequency analyses will provide changes in normalised PSD in units of standard deviation (SD).

### Statistical analysis

To test our hypotheses, we conducted several statistical analyses including within-subject nonparametric permutation tests and circular statistics. All statistical analyses were performed using custom scripts in Matlab (v. 2022b), Python (v. 3.9.7), and R (v. 4.2.2).

Timewise permutation tests were carried out on ECG and EMG data, to assess within-subject differences as described in “*EEG and EMG Data Analysis*”. In the EEG data, we conducted timewise permutation tests to compare the normalised PSD of alpha and beta between ipsilateral and contralateral traces. This analysis was carried out first on the entire dataset including all trials regardless of cardiac information and then separately on systole and diastole trials. The primary analysis focused on data source-reconstructed within the 500–2000 ms time window, with averaging in the alpha (8–13 Hz) and beta (14–30 Hz) frequency ranges to yield one frequency bin per band. Additionally, as an exploratory step, the analysis was extended to cover the full epoch length, from 500 to 4000 ms post-cue. For all permutation tests, we used 500 permutations and set an alpha level of 0.05 (Marozzi, 2004). For two-sided permutation tests, we considered statistical values that fell in the left (2.5%) and right (97.5%) percentile of the permutation distribution (two-tailed, *p* < 0.025). In cases of multiple comparisons, we controlled the false discovery rate (FDR) at *q* = 0.05 using an adaptive linear step-up procedure (Benjamini, Krieger, & Yekutieli, 2006), and reported the significant results using the adapted threshold *p*-value (*P_FDR_*). Non-significant effects after FDR control are presented simply by the p-value (*P*). Nonparametric effect sizes were estimated using the probability of superiority for dependent samples (Δ_dep_) (Grissom & Kim, 2011; Ruscio & Mullen, 2012).

The EEG analysis was complemented with a control analysis to account for the varying lengths of the systole and diastole phases. Due to the longer duration of the diastole phase, more trials occurred during diastole than during systole. To address the potential effect of this imbalance on alpha and beta PSD changes, we randomly selected an equal number of trials from both the systole and diastole phases across the entire dataset. This process of random selection (with replacement) was repeated ten times—control runs. For each run, we conducted the same timewise permutation tests, applied the adaptive FDR control procedure, and estimated nonparametric effect sizes.

Next, complementing the primary EEG, ECG, and IBI analyses, we investigated the (i) temporal alignment of the experimental cue with the cardiac cycle, to assess whether participants aligned their heartbeats to the cue presentation. In addition, we examined (ii) the alignment of cue latency in MI trials that exhibited more pronounced contralateral suppression in alpha and beta frequencies, relative to ipsilateral changes, using a predefined suppression threshold (see below). In both instances, we examined the specific angular positions of the cues within the cardiac cycle, analysing continuous angles from 0 to 2π radians (*rad)*, instead of categorising them into binary phases like diastole or systole. These analyses employed the Rayleigh test, which is suitable for angular data with periodic oscillations, such as cardiac activity. This test determines whether the distribution of data around a circle is non-uniform, indicating concentration at specific angles.

For the temporal alignment analysis (i), we converted the latency difference between the cue and the preceding R-peak into angles (θ) in radians. This conversion was done after normalising the latency difference with the IBIs from the last four R-peaks, using the following expression:

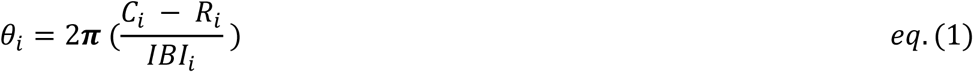

where

- *C*_*i*_ = The onset of the cue.
- *R*_*i*_ = The onset of the R-peak before the cue of interest.
- *IBI*_*i*_ = The average cardiac cycle obtained from the previous four cycles.
- *i* = The i^th^ trial.

In the EEG analysis (ii), we computed a trial-wise suppression index (SI) by subtracting the normalised alpha and beta PSD values of the contralateral side from those of the ipsilateral side for each trial (SI: trial-wise ipsilateral - contralateral PSD). These differences were then averaged over a significant post-cue interval, identified from the primary contralateral versus ipsilateral statistical contrasts conducted independently of cardiac effects (i.e. using all trials to avoid bias in circular statistics). A positive trial-wise SI indicates relative contralateral suppression, whereas a negative SI implies ipsilateral suppression. To determine if there is an optimal cue presentation timing for enhancing contralateral alpha and beta suppression during MI, which could be useful to guide follow-up BCI work, we analysed the angles (θ) of trials exceeding the 50th, 75th, and 90th percentile thresholds for contralateral suppression, after converting them into radians. Thresholds were estimated in each participant. The Rayleigh test was then applied to these angular data for alpha and beta frequency bands, as well as for the M1 and S1 cortices, separately. Initial tests were conducted at the individual level to derive *p-values* and resultant vector values for single-subject effects. This was followed by a group-level analysis, using the individual mean resultant vectors, to identify any directional bias in the circular distribution of suppression-related cues on the group level. Statistical effects are provided with uncorrected *p-values* as this analysis served to explore potential time windows of clustering relevant to guide future real-time BCI studies.

Last, to determine whether the directionality effects (circular statistics) of the SI analysis along the cardiac cycle could be explained by randomly distributed values along the unit circle, we created a null distribution in each subject, by randomly generating angles from 0 to 2π for each trial and subject, representing random cue latencies of above-threshold power suppression values. Single-subject Rayleigh tests were followed by group-level Rayleigh tests. This analysis helped determine whether the circular distribution of randomly generated cue latencies could explain significant group-level effects, as we hypothesised for our experimental PSD suppression data.

## Results

### Assessing differences between ECG and EMG waveforms

In the ECG data, within-subject nonparametric permutation tests found no significant differences in the waveforms between the left-cued and right-cued trials for both MI and ME tasks (*P* > 0.05 after FDR control; see **Figure 2a)**. Additionally, there were no significant differences between participants’ IBIs in both motor tasks (*P* > 0.05 after FDR control; **Figure 2b**).

**Figure 2.**
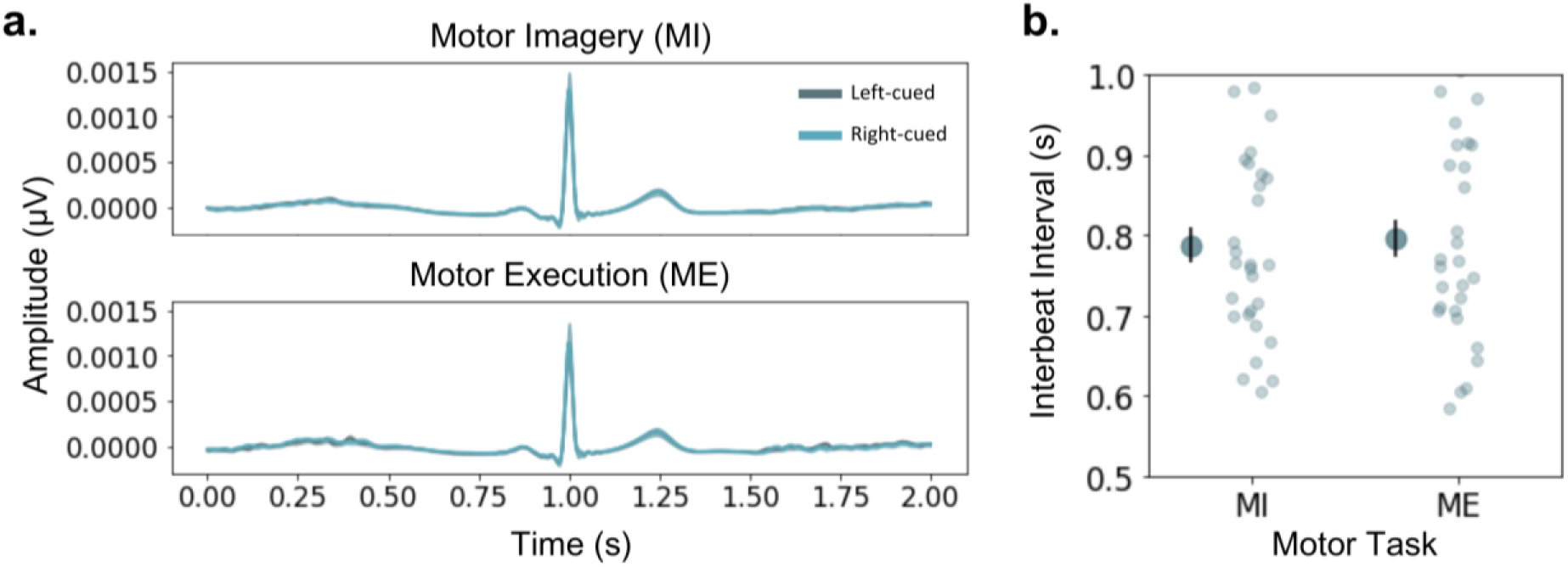
Differences between ECG waveforms and IBIs. **a.** The left panel shows the overlap between the ECG waveforms of left-cued (dark blue) and right-cued (light blue) in the MI (top) and ME (bottom) tasks. **b.** The right panel compares the IBIs between MI and ME. The larger darker dots are the mean, with SEM represented by the vertical black bar. Lighter smaller dots are the individual data points for each participant. For both panels, non-significant differences were observed between the ECG waveforms across conditions (**a**) and participants’ IBIs (**b**).

Subsequently, we examined EMG differences between the ipsilateral and contralateral traces in MI and ME tasks (**Figure 3**). We observed a significant difference in the ipsilateral minus contralateral activation during the ME (*P_FDR_*=0.0001–0.0108, range of *p-values* for time points associated with significant differences after FDR control*, Δ_dep_*=0.8846) and, unexpectedly, MI task (*P_FDR_*=0.0001*, Δ_dep_*=1). The significant effects extended for the entire window (500–2000 ms), as shown in **Figures 3a** and **3b**, respectively. Although the significant lateralisation of EMG activity during motor imagery was unexpected, indicating residual lateralised muscle activation in participants during MI, we confirmed in a post-hoc analysis that EMG activity during ME was significantly greater than during MI (*P_FDR_*=0.0001*, Δ_dep_*=1).

**Figure 3.**
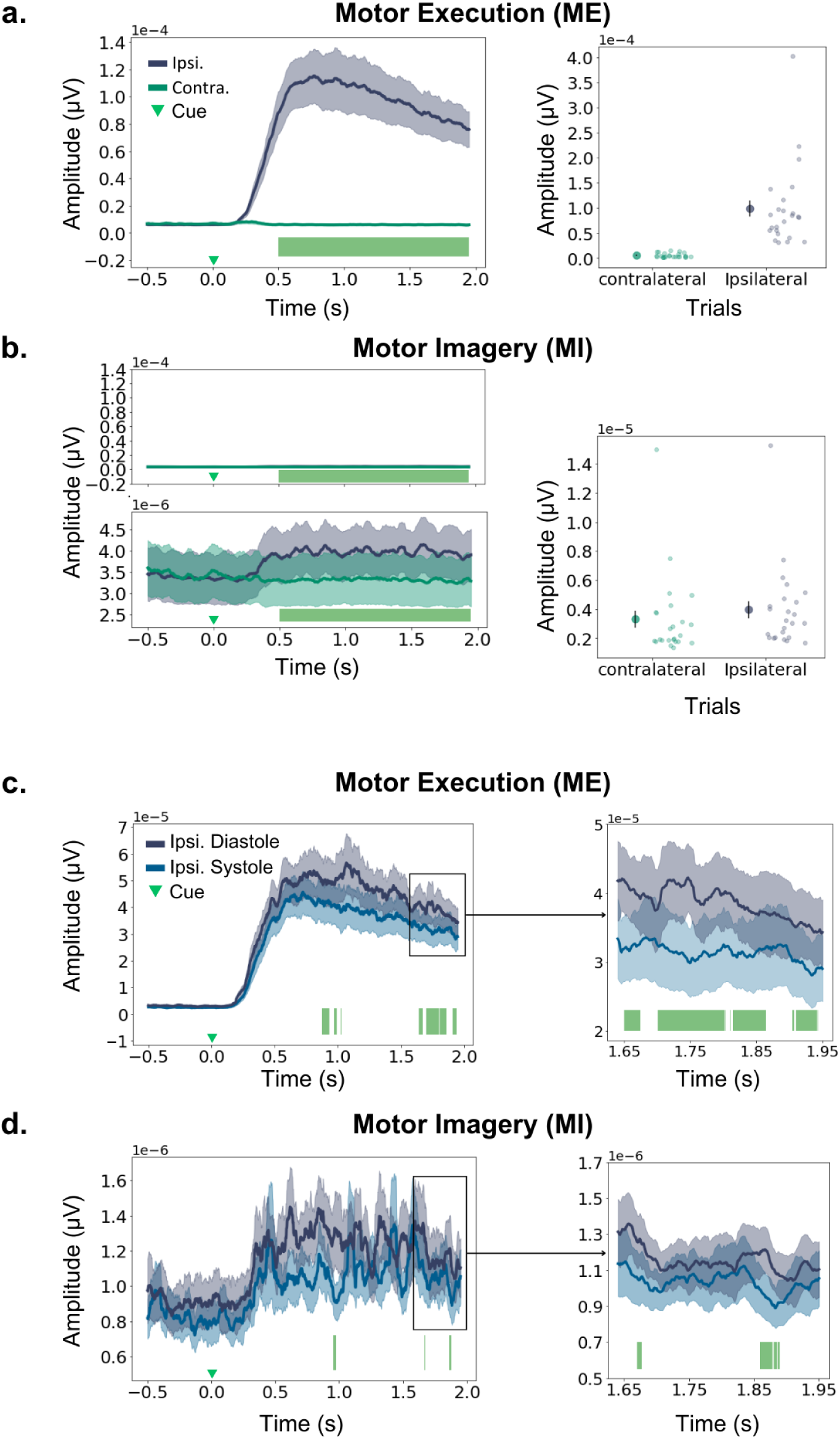
Differences between EMG waveforms. **a-b. Left.** Time course of ipsilateral and contralateral EMG waveforms during ME (**a**) and MI (**b**) tasks. Non-parametric permutation tests assessing EMG differences between ipsilateral and contralateral traces revealed a significant increase in ipsilateral muscle activity, relative to contralateral, for the ME (**a**) and MI (**b**) tasks. *P-values* for significant effects after FDR control were in the range *P_FDR_=*0.0001–0.0108 for ME, *P_FDR_=*0.0001, for MI. Significant effects are denoted by the green bars at the bottom. This test was conducted in the main window of interest, up until 2 s after the cue. Panel (**b**) includes two subpanels: The top panel uses the same y-axis scale as in ME (**a**) to allow for a qualitative visualisation of the difference in muscle activity between the two tasks. The bottom panel adjusts the y-axis scale to the data, to illustrate the laterality modulation. **Right**. Average EMG amplitude for each participant (lighter dots) and the group mean (darker dot) obtained from contralateral and ipsilateral hands (SEM is shown with the vertical black bar). All statistical analyses were conducted in the subsample where EMG was available (n=26), however, for illustration purposes, two participants are not included in this graphic as they had very large amplitude EMG values. Therefore, Figure 3 represents data in n=24 participants. The panels including all 26 participants can be found in **Figure S1**. We did not exclude these two participants with larger EMG amplitude modulations from statistical analyses because the changes were of the same latency as in the other participants, and larger values may have been simply due to larger SNR on the day of the recording. **(c-d)**. Same as (a) but comparing ipsilateral EMG traces when cues instructing movement direction occur in the diastolic (darker blue) or systolic (lighter blue) phase of the cardiac cycle. Significant effects are denoted by the green bars at the bottom, with *p-values* in the range *P_FDR_=*0.0001–0.0134 for ME, *P_FDR_=*0.0001–0.002, for MI. **Right**: The right-side panels represent zoomed-in inserts of the EMG activity between 1.65 and 1.95 s. For all waveforms, SEM above and below the group average is shown in lighter colour areas. See **Figure S2** including all n=26 participants with available EMG data. As in (**a-b**), all statistical analyses represented in panels (**c-d**) included all n=26 participants, but the EMG traces represent the subset of n=24 for clarity. See main text.

Additionally, based on recent findings of TMS-related EMG differences as a function of the cardiac cycle (Al et al 2023, but see Bianchini et al. (2021) for contrasting results), we carried out an analysis to determine the difference between the EMG activation (ipsilateral traces) cued during systole and diastole. We found a significant difference in ME (*P_FDR_=*0.0001–0.0134, *Δ_dep_*=0.92; see), but also in MI (*P_FDR_=*0.0001–0.002, *Δ_dep_*=0.81), due to larger EMG activity in diastole. The latency of the effects showed partial overlap for both tasks, occurring at approximately 0.87–1.02 s and 1.64–1.94 s during ME (**Figure 3c**), and at approximately 0.95–0.97 s, around 1.67 s, and 1.85–1.88 s during MI (**Figure 3d**). However, the significant effects during MI trials lasted only a few milliseconds.

### Source Analysis: Systole vs Diastole

When participants overtly executed thumb abduction, our analysis did not detect a significant lateralisation effect in the normalised PSD within the alpha and beta frequency ranges for both the M1 and S1 cortical regions. This absence of significant effects manifested when using all trials, as well as in the datasets of trials cued during systole and diastole (P > 0.05 after FDR control in all cases). No significant effects were observed in the exploratory analysis considering the full epoch length (0.5–4.0 seconds) either.

During MI, we observed a significant lateralisation of alpha and beta suppression in the dataset including all trials (**Figure 4a**). This was due to more pronounced contralateral suppression in both the alpha (S1: *P_FDR_*=0.0001–0.0006*, Δ_dep_*=0.79; no effect in M1: *P* > 0.05 after FDR control) and beta bands (M1: *P_FDR_*=0.0001–0.0006*, Δ_dep_*=0.79; S1: *P_FDR_*=0.0002– 0.001*, Δ_dep_*=0.76). In both regions, these significant effects were short-lived, extending for approximately 60 ms in alpha (S1: 1531–1589 ms) and 20 ms in beta (S1: 582–605 ms; M1: 500–523 ms), and therefore did not cover a full cycle at the central frequency in those bands (100 ms for 10 Hz, 50 ms for 20 Hz).

**Figure 4.**
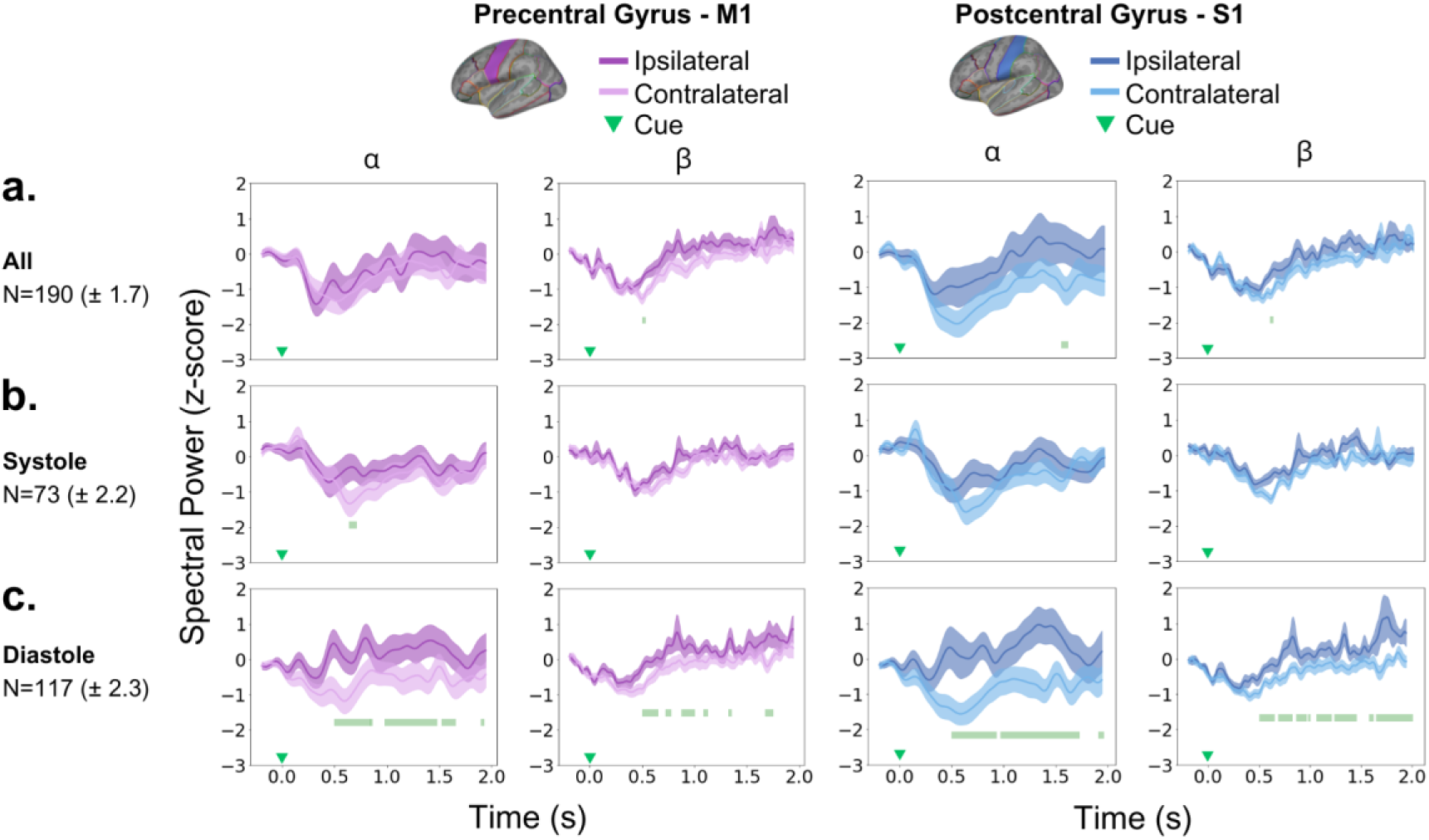
Differences between Ipsilateral and Contralateral Power Spectral Density in M1 and S1. **a-c.** Normalised PSD in alpha and beta bands, separately in the precentral gyrus (M1, purple traces) and postcentral gyrus (S1, blue traces). Panel **(a)** shows the time points associated with significant differences between ipsilateral (darker colour) and contralateral (lighter colour) traces for all trials (N = 190 ± 1.7), denoted by the green bars at the bottom. Continuous lines represent the group-average, while shaded areas represent the SEM. For all trials, a significantly larger suppression was found for alpha in S1 (top-left panel, *P_FDR_=*0.0001–0.0006), and for beta in M1 (top-left panel, *P_FDR_=*0.0001–0.0006) and S1 (top-right panel, *P_FDR_=*0.0002–0.001). (**b-c**). Same as (**a**) but for systole-cued trials and diastole-cued trials, respectively. Contralateral suppression was sustained in diastole-cued trials for alpha (bottom-left, M1: *P_FDR_=*0.0001–0.0544; bottom-right, S1: *P_FDR_=*0.0001–0.0592) and beta (bottom-left, M1: *P_FDR_=*0.0001–0.0578; bottom-right, S1: *P_FDR_=*0.0001–0.0576) bands. In systole-cued trials, a short-lived effect was observed only in the alpha-band PSD of M1 (middle-left panel, *P_FDR_=0*.0002–0.0004).

Upon separately analysing laterality effects for trials cued during systole and diastole, in the systole-cued trial subset (**Figure 4b**), significant differences were also confined to a brief time window (640–700 ms), exclusively in the alpha band and M1 region (*P_FDR_*=0.0002– 0.0004*, Δ_dep_*=0.79). No additional significant effects were observed for systole-cued trials (alpha, S1: *P* > 0.05 after FDR control; beta, S1 and M1: *P* > 0.05 after FDR control). Conversely, pronounced contralateral suppression effects were observed in diastole-cued trials (**Figure 4c**), marked by significant differences in the alpha (M1: *P_FDR_*=0.0001–0.0544*, Δ_dep_*=0.83; S1: *P_FDR_*=0.0001–0.0592*, Δ_dep_*=0.79) and beta bands (M1: *P_FDR_*=0.0001–0.0578*, Δ_dep_*=0.86; S1: *P_FDR_*=0.0001–0.0576, *Δ_dep_*=0.83). These effects persisted throughout the entire time window, starting at 500 ms and continuing until approximately 1800–1900 ms after the onset of the S1 (alpha: 500–1931 ms; beta: 500–1953 ms) and M1 (alpha: 500–1918 ms; beta: 500–1742 ms).

Exploring potential lateralisation effects over the entire epoch (0.5–4.0 seconds) yielded similar findings, as shown in **Figure S3**. In the total dataset, significant but brief contralateral suppression occurred in both alpha (S1: P_FDR_=0.0004–0.0012, Δ_dep_=0.72) and beta (M1: P_FDR_=0.0001–0.0012, Δ_dep_=0.79; S1: P_FDR_=0.0004–0.0012, Δ_dep_=0.83) bands, with effects lasting approximately 100 ms and 10 ms at different times. Notably, in diastole-cued trials, significant contralateral suppression was observed in alpha (M1: P_FDR_=0.0044–0.0094, Δ_dep_=0.76; S1: P_FDR_=0.0001–0.01, Δ_dep_=0.9) and beta (M1: P_FDR_=0.0001–0.0094, Δ_dep_=0.86; S1: P_FDR_=0.0001–0.01, Δ_dep_=0.93) frequencies. No significant differences were obtained in systole-cued trials.

To determine if participants’ heartbeats aligned with the cue presentation, we carried out the Rayleigh test on angular data representing the latencies between the cue and the preceding R-peak. The results were not significant, suggesting a lack of effects for a deviation from a uniform distribution around the circle (see **Figure S4**).

After balancing the number of trials for systole-cued, diastole-cued, and the full dataset across 10 control runs, the main effects remained consistent, as illustrated in **Figure 5**. This confirmed that the observed significant and sustained contralateral suppression in alpha and beta PSD within sensorimotor cortices during diastole-cued trials was not due to a higher trial count compared to systole-cued trials. Refer to **Tables S1–4** for the corresponding statistics. Indeed, even when considering all trials, systole-cued and diastole-cued, the effect was less sustained, as reported above (**Figure 4**).

**Figure 5.**
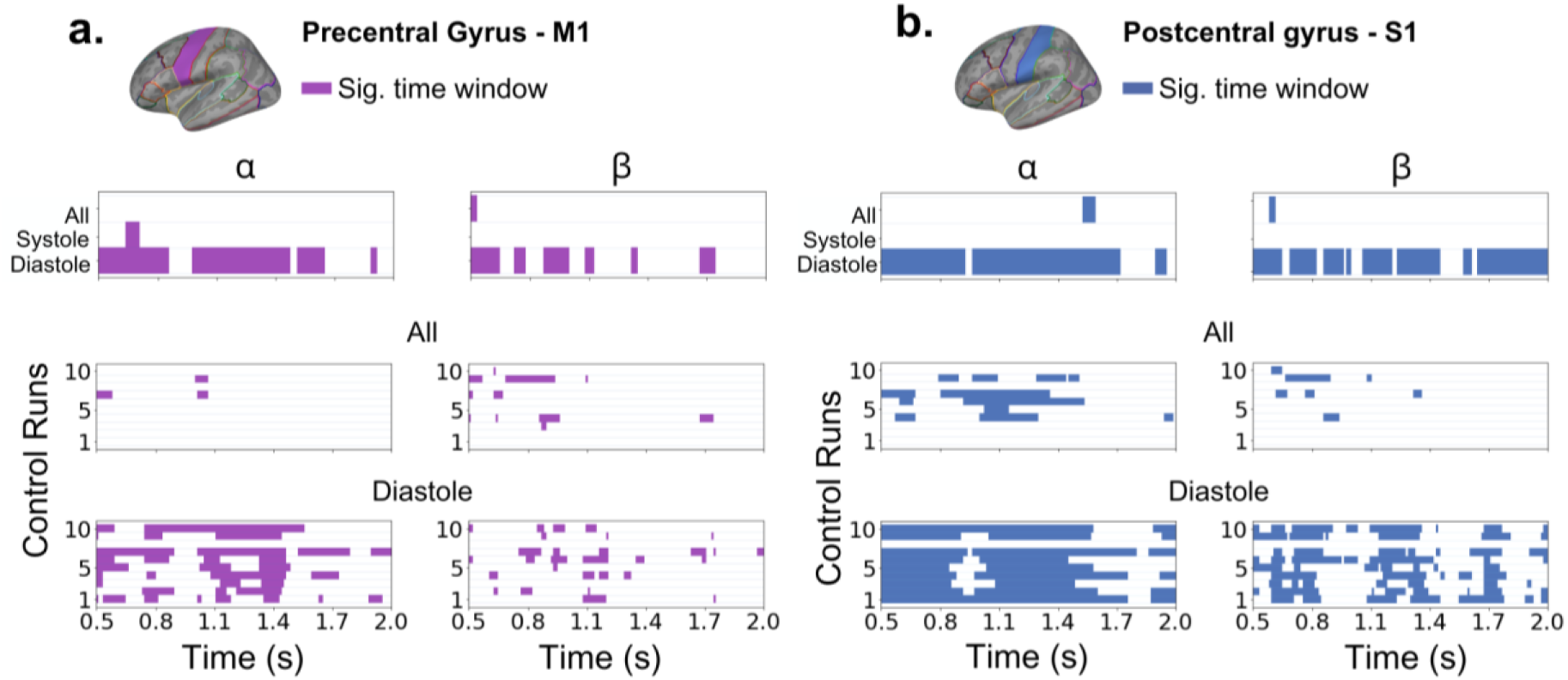
Control analysis of laterality effects by cardiac phase. The figure shows the results of the control analysis (10 runs) in the M1 (**a**) and S1 (**b**) cortical regions. The coloured horizontal bars denote time windows of significant differences between contralateral and ipsilateral suppression after FDR control, and M1 (purple) and S1 (blue) regions. Top panels summarise the statistical effects of the main analysis, including all available trials of each type: irrespective of cardiac cycle phases (“All”), systole-cued trials, diastole-cued trials. The middle panels illustrate statistical effects in the 10 control analyses conducted using a reduced number of trials, matched to the number of systole-cued trials for each participant. This revealed a few laterality effects for M1 (alpha: 2/10 runs; beta: 6/10) and S1 (alpha: 5/10; beta: 4/10), albeit in time points not overlapping with the windows showing effects in the main analysis (top panel). See range of *p-values* and effect sizes in **Tables S1-S4**. The bottom panels show the time intervals associated with significant laterality effects, obtained using the reduced number of diastole-cued trials. Significant effects were widespread and consistent across runs, in line with the main analysis: significant effects observed for M1 (alpha: 9/10 runs; beta: 8/10) and S1 (alpha: 9/10; beta: 9/10).

Last, although a robust and widespread significant lateralisation effect of alpha and beta suppression in diastole-cued trials during MI was observed with cues aligned with diastole, we conducted a visual inspection to determine if the onset of significant effects coincided with the systole phase. This post-hoc analysis was motivated by recent work indicating heightened motor cortex excitability when TMS is administered during systole. As shown in **Figure 6**, when cues were initiated during diastole, the onset of significant suppression effects did not consistently align with the systole phase across participants. Furthermore, the significant effects lasted from 0.5 to 1.9 seconds, indicating that the enhanced contralateral suppression observed in diastole-cued trials extends across various phases of the cardiac cycle, transitioning from diastole to systole.

**Figure 6.**
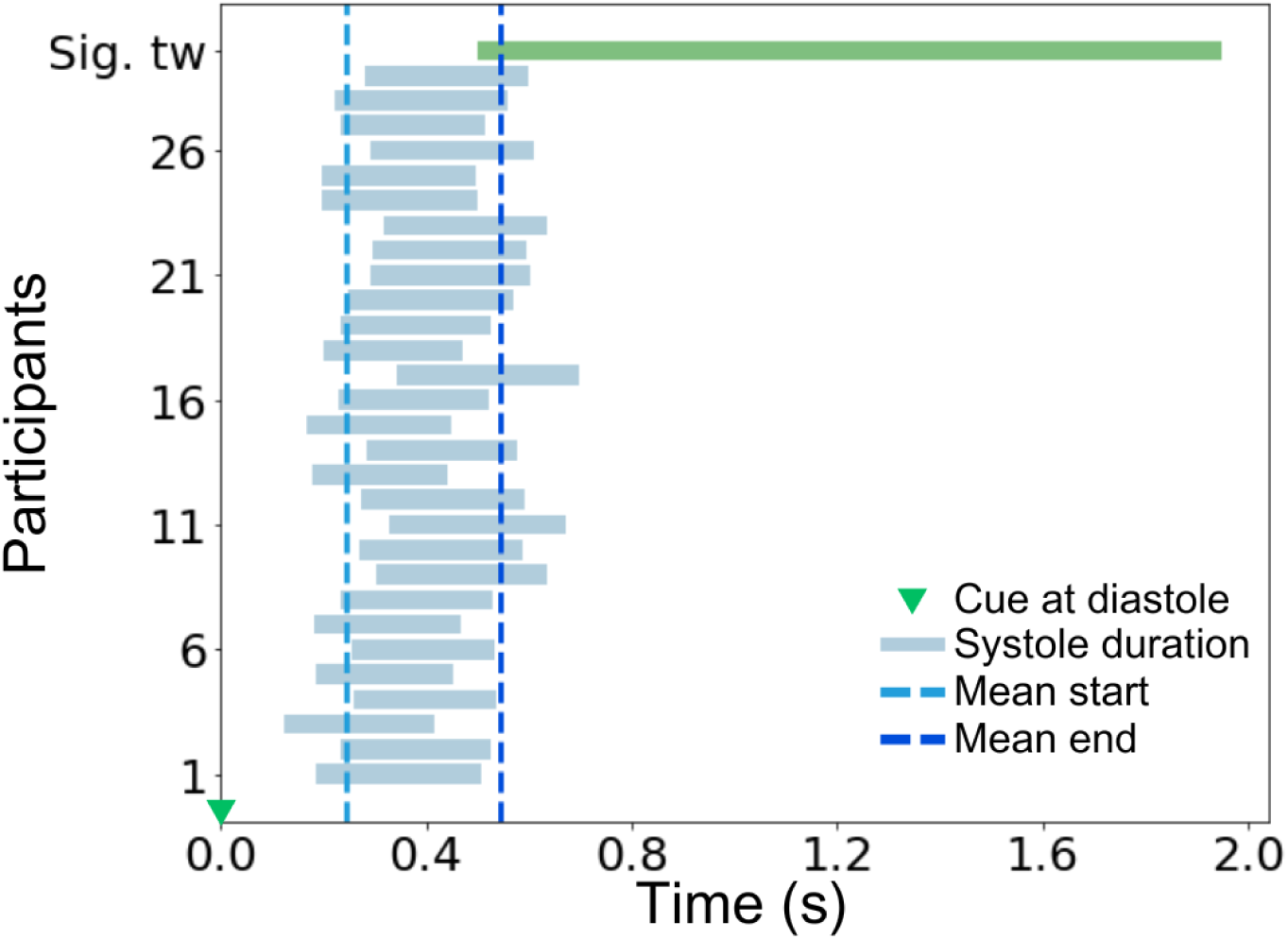
Latency of significant lateralisation effect of alpha and beta suppression in diastole-cued trials. The onset of significant lateralisation effects in S1 and M1 around 0.5 s (extending during 500–1900 ms, denoted by the green bar on top) did not consistently overlap with the duration of the subsequent systolic phase (mean duration: 243–546 ms) for each participant. On average, the overlap was for 46 ms only. Time zero coincides with the presentation of the cue during the diastolic phase.

### Position of the cue within the cardiac cycle and contralateral suppression

We employed circular statistics to evaluate whether the latency of cues in trials with particularly pronounced contralateral alpha and/or beta suppression clustered at a specific angle on the unit circle, representing the cardiac cycle. This process involved initially calculating the contralateral suppression index (SI: PSD ipsilateral - contralateral) for each trial and participant. Positive SI values indicate more contralateral suppression, while negative values denote greater ipsilateral suppression. To identify trials with “pronounced suppression” we applied a criterion based on three different data embeddings: selecting trials that fell at the 50th, 75th, or 90th percentile of the SI distribution across all trials separately for each participant. First, at the individual participant level, the Rayleigh test did not reveal any significant effects across most participants. However, at the group level, a significant departure from a uniform distribution was observed in the M1 region for SI values exceeding the 50th percentile, associated with trials of “particularly pronounced” contralateral suppression in the alpha frequency range (Rayleigh’s Z=0.3229, P=0.04727; **Figure 7ab**). The direction of the group-level mean resultant vector was 4.59 *rad*, within the diastole. In the S1 region, a significant deviation was only detected for contralateral alpha suppression above the 75th percentile (Rayleigh’s Z=0.3298, *P*=0.0412; **Figure 7de**). This effect was associated with a mean resultant vector aligned at 3.84 *rad*, also placed within the diastole phase. Other combinations of percentile threshold embedding, ROI and frequency band did not yield significant results.

**Figure 7.**
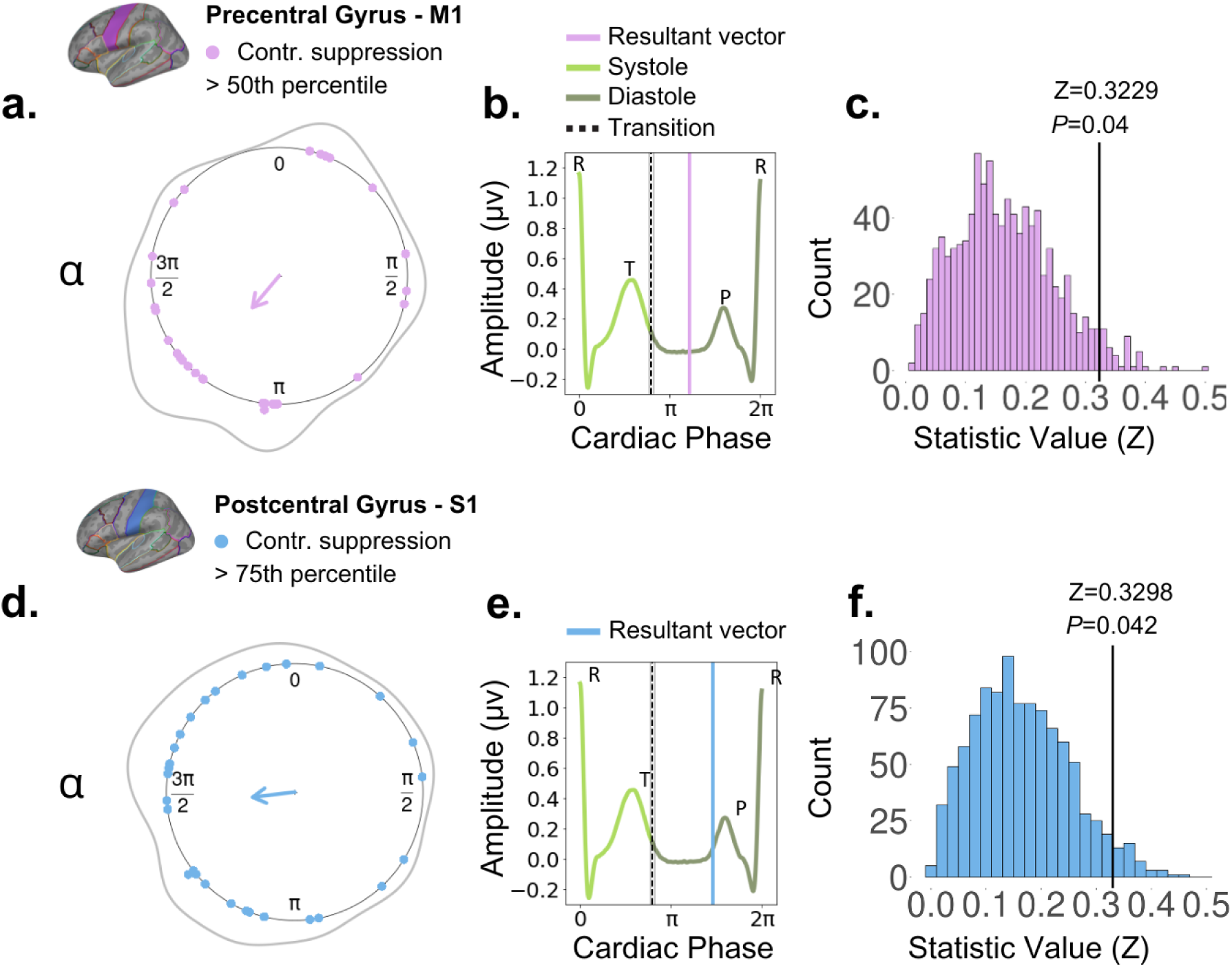
Temporal clustering along the cardiac cycle for cues associated with trials exhibiting contralateral suppression. We used circular statistics to assess whether the latency of cues in trials in which there was contralateral alpha and/or beta suppression clustered at a particular angle along the unit circle, representing the cardiac cycle. The top and bottom panels refer to alpha in the 50^th^ and 75^th^ percentiles, respectively. From left (**a, d**), the circular plots show the mean resultant vector (group-level statistics) pointing at the angle in which clustering occurred. Individual dots represent the subject-specific direction of the mean resultant vector. The Rayleigh test was significant in M1 when using a 50th percentile threshold (**a**; Z=0.3229, *P*=0.04727), and in S1 when using a 75th percentile threshold (**d**; Z=0.3298, *P*=0.0412). In both cases, this indicates a deviation from the uniform distribution. In the middle panels (**b, e**), we show the position of the angle of the group-level mean resultant vector with respect to the ventricular diastole (vertical purple line: top, blue line: bottom). The dashed line with shaded areas represents the group-average transition from systole to diastole (end of the T-wave). The right-side panels (**c, f**) present the empirical Rayleigh *Z* statistic compared to a null distribution of *Z* statistical values. These values are estimated from 1,000 Monte-Carlo-based permutations, which consist of randomly generated angles for each trial and participant. This approach involves subject-level analysis, subsequently followed by group-level analysis.

In a control analysis, we compared the empirical Rayleigh Z statistic corresponding to the group-level significant SI effects to a null distribution of Z statistical values based on 1,000 Monte-Carlo-based permutations. This analysis revealed a significant effect in both cases (*P*=0.04 for M1 and 50th percentile, *P*=0.042 for S1 and 75th percentile). That means that the clustering of trials of particularly pronounced alpha or beta suppression at a specific angle during diastole was not explained by a random distribution of trial-related and subject-related angles on the unit circle.

## Discussion

Utilising a motor imagery task, where participants were instructed to mentally imagine the kinesthetic sensations of lifting their left and right thumbs, we found that the timing of the cardiac cycle modulates the contralateral alpha and beta suppression in sensorimotor cortices. This is the first study to demonstrate that MI performance—indexed by enhanced alpha and beta PSD suppression over the contralateral sensorimotor cortex—is significantly improved when the experimental cue instructing movement direction coincides with the diastolic phase, rather than the systolic phase. These results are in line with the prominent baroreceptor hypothesis (Lacey, 1967), which proposes that heightened baroreceptor activation during systole might lead to a diminished response to sensory cues occurring in this systolic phase. We discuss these findings and their potential application for the development of assistive and rehabilitative technologies, including methodological implementations that could improve MI-based BCIs and neurorehabilitation training, especially for post-stroke patients.

### Contralateral Alpha and Beta Suppression during Motor Imagery

When analysing neural responses to imagined movements, it is expected that imagining kinesthetic sensations leads to desynchronisation in the hemisphere contralateral to the imagined movement (Neuper, Scherer, Reiner, & Pfurtscheller, 2004). We observed significantly more pronounced suppression of alpha and beta activity in the S1 and M1 regions of the contralateral hemisphere during the MI task. Notably, in S1, the significant alpha desynchronisation started at approximately 1500 ms post-cue and lasted less than 100 ms, with no effects in M1. Similarly, suppression in the beta band was transient in both S1 and M1, appearing around 580 or 500 ms, respectively, and lasting less than 50 ms. Despite the transient nature of the significant effects, the findings align with the literature (Grèzes & Decety, 2001, Hetu et al., 2013; Porro et al., 2000), indicating that imagining the sensation of movement activates cortical networks within S1 and M1.

However, during actual motor execution, we did not find significant lateralisation effects. Such absence of larger contralateral modulation during real motor performance is not uncommon. A recent review supports the notion that unilateral movements, such as wrist extension or thumb abduction as in our study, lead to bilateral cortical activation, with lateralisation in such cases seeming to emerge with higher motor complexity (Chettouf, Rueda-Delgado, de Vries, Rittera, & Daffertshofer, 2022; Li et al., 2018). The authors have further concluded that this phenomenon could potentially be due to neural activation patterns involving both inhibition and excitation across the bilateral motor cortex during unilateral movements, with transcallosal tracts facilitating synchronous communication between hemispheres.

### Cardiac Cycle Modulation

When assessing our hypothesis that the cardiac cycle modulates neural responses during MI, we found that alpha and beta contralateral suppression in both M1 and S1 regions was more pronounced when the experimental cue occurred in the diastolic phase, with almost no effects for systole-cued trials. Specifically, effects in diastole-cued trials were more sustained than those observed across the entire dataset, lasting the full duration of the analysed time window (0.5–2 seconds) in both ROIs. These findings align with the baroreceptor hypothesis (Lacey, 1967), which implies that during diastole, when the activation of mechanoreceptors in cardiac walls does not attenuate the signal-to-noise ratio (unlike during systole), neural and behavioural responses across different sensory modalities can be enhanced (Azzalini et al., 2019; Engelen et al., 2023). Conversely, during systole, there was a minimal lateralisation effect in M1 that was short-lived, lasting only 60 ms. These results offer preliminary evidence that MI performance can be more effectively modulated within the context of the cardiac cycle.

Our post-hoc analyses, aimed at exploring cardiac modulation of contralateral alpha and beta PSD suppression over a prolonged time window (0.5–4 seconds), confirmed the main analyses, revealing that the significant modulatory effects in S1 and M1 in diastole-cued trials were predominantly confined within the first two seconds across the entire dataset. We additionally addressed potential concerns about the imbalance in the number of trials between diastole and systole phases, as highlighted in previous research (Boudewyn, Luck, Farrens, & Kappenman, 2019), by matching the number of trials in diastole to those in systole at the subject level. Our control analysis showed that the effects in sensorimotor cortices, and particularly for S1, are robust even with a reduced number of trials. Interestingly, this robustness of the diastole effects did not extend to the full trial dataset analysis, as reducing the number of trials— therefore containing a random subset of diastole and systole trials—led to a reduced number of significant results.

Finally, using circular statistics, we demonstrated that trials with particularly pronounced contralateral suppression tended to cluster at specific times during the diastole phase. Specifically, in M1, large contralateral suppression in alpha clustered approximately 430 ms after the T-wave offset (50^th^ percentile thresholding), while in S1 clustered around 270 ms (75^th^ percentile thresholding). Both timing effects coincide with the lowest cardiovascular arousal period during the diastolic phase (Kurbel, 2004), between the T-wave (ventricular repolarisation) and the P-wave (atrial repolarisation). In the ECG, the moment between these noisy events is known as the isoelectric line or baseline (Kurbel, 2004), which is considered to be the rest phase before a new cardiac cycle begins.

Interestingly, our findings in the ME domain do not support the notion that higher cardiovascular arousal enhances neural responses during overt movements, as observed in a recent study by Al et al. (2023). In their TMS study, Al and colleagues demonstrated that finger movements triggered during systole, led to larger MEPs and more pronounced desynchronisation in the alpha and beta frequency ranges. First, our results show lack of lateralised suppression when ME trials were executed during systole and diastole, similarly to what we found when all trials were considered. Moreover, assessing the differences between ipsilateral EMG traces of movements cued during systole and diastole revealed significantly higher muscle activity for the diastolic phase, which occurred at approximately 900 ms and continued later between ∼ 1650-1950 ms. There is, however, an important methodological difference that needs to be taken into account. In Al and colleagues’ study, finger movements were involuntary and triggered by TMS, meaning that the muscle response was relatively instantaneous from receiving the pulse. In our study, muscle response was delayed by approximately 500 ms post cue (ipsilateral versus contralateral EMG analysis), and initiated by the participants, as shown in Figure 4, rendering both sets of results not directly comparable.

Additionally, upon visualising where the onset of significant contralateral alpha and beta suppression effects in diastole-cued trials occurred, we noted minimal overlap with the systole phase across participants. Moreover, the neural effects lasted for up to 2 seconds, complicating any inference about the effect of the cardiac cycle on the movement itself. Overall, our findings indicate that enhanced sensory processing, occurring when the movement cue is presented during the diastole phase, might better predispose participants to execute the imaginary movement at a later point, post-diastole cycle, as indicated by alpha and beta PSD suppression.

### Applications

Our findings have practical implications for assistive technologies, especially MI-based BCIs for individuals with severe neurological disorders and MI-based neurorehabilitation for post-stroke upper limb recovery. In MI-based BCI training, users generate MI examples at specific times following system prompts, which is crucial for collecting labelled data to train supervised machine learning models (Sannelli et al., 2019). These models decode users’ intentions in real-time during the BCIs’ communication phase (Sannelli et al., 2019). Our results suggest that instructing users to generate MI examples during diastole, rather than systole, during training could increase the likelihood of capturing contralateral sensorimotor suppression. This should be assessed in follow-up work. Additionally, the potential enhancement of kinesthetic sensations from diastole-cued movements requires further validation in future research.

In neurorehabilitation, MI training has shown significant benefits in aiding stroke survivors regain motor function in affected limbs. For example, Daly and Bialocerkowski (2009) found MI training enhanced motor control by 80%. Studies have also highlighted short-term advantages of MI for reducing motor spasticity, increasing muscle strength, improving daily activities, and facilitating long-term muscle recovery (Biasiucci et al., 2018; Pichiorri et al., 2015; Ramos-Murguialday et al., 2013; Bai et al., 2020). Our findings suggest the potential for more dynamic neurorehabilitation protocols that consider the heart-brain interaction.

Encouraging MI exercises during diastole could improve performance and kinesthetic sensations and might foster neural plasticity. This approach, which should be first tested in healthy individuals with real-time cardiac phase monitoring, could extend to patient groups using offline analysis to monitor rehabilitation progress. Integrating recent advances in real-time ECG analysis for assessing cognitive-emotional control (Adelhöfer et al., 2020) into this framework could also help refine cardiac-cycle dependent neurorehabilitation and BCI protocols further.

### Limitations

Our study is not without its limitations. First, we suggest that triggering MI trials during diastole may increase the engagement of the contralateral sensorimotor cortex and potentially improve MI performance through better kinesthetic sensations. However, we did not collect data on participants’ kinesthetic experiences. Future studies should aim to replicate these findings while employing methods to quantify vividness and kinesthetic sensations during MI, such as the Kinesthetic and Visual Imagery Questionnaire (KVIQ; Malouin et al., 2007; Yang, Jeon, Kim, & Chung, 2021). Incorporating such measures could elucidate the relationship between MI performance during different phases of the cardiac cycle and the vividness and sensation scores.

Second, for our results to be fully applicable in MI-based BCIs, they should also be replicated in an extended version of the current experimental paradigm that involves multiple sessions over several days and incorporates visual feedback to guide the learning process. In previous studies, multiple sessions and visual feedback demonstrated more concentrated desynchronised neural responses in the motor cortex (Rimbert & Fleck, 2023) and promoted learning by enhancing a sense of agency and embodiment towards the task (Alimardani, Nishio, & Ishiguro, 2018). Future studies could aim to combine these two factors to gain a deeper understanding of how the cardiac cycle influences MI performance.

### Conclusion

Our study highlights the significant influence of the cardiac cycle on MI performance. We found that periods of lower cardiovascular arousal, between the T-wave and the P-wave of the diastolic phase, may enhance the processing of experimental instructions (left/right arrows) used to guide imaginary movements. While further studies are needed to ascertain if this advantage during diastole can be detected in real-time, and whether it can be generalised to other imaginary tasks, our findings contribute valuable insights into the impact of the cardiac cycle on the variability of MI performance, with promising implications to improve MI-based assistive technologies.

## Author contributions

G.L. contributed to conceptualisation, investigation, methodology, software, formal analysis, visualisation, writing-original draft preparation, funding acquisition. M.H.R. contributed to conceptualisation, methodology, software, formal analysis, visualisation, writing-reviewing and editing, supervision, funding acquisition. D.L. contributed to conceptualisation, funding acquisition. C.V. and J.B. contributed to writing-reviewing and editing.

## Acknowledgments

The study was supported by Liquidweb s.r.l. and Goldsmiths, University of London, funded through the industry-sponsored project “Using heart-brain interactions in brain-computer interface systems” ID 408252.

## References

Adelhöfer, N., Schreiter, M. L., & Beste, C. (2020). Cardiac cycle gated cognitive-emotional control in superior frontal cortices. NeuroImage, 222, 117275. 10.1016/j.neuroimage.2020.117275

Al, E., Stephani, T., Engelhardt, M., Haegens, S., Villringer, A., & Nikulin, V. V. (2023). Cardiac activity impacts cortical motor excitability. PLOS Biology, 21(11), e3002393. 10.1371/JOURNAL.PBIO.3002393

Alimardani, M., Nishio, S., Ishiguro, H., Alimardani, M., Nishio, S., & Ishiguro, H. (2018). Brain-Computer Interface and Motor Imagery Training: The Role of Visual Feedback and Embodiment. Evolving BCI Therapy - Engaging Brain State Dynamics. 10.5772/INTECHOPEN.78695

Arnau, S., Sharifian, F., Wascher, E., & Larra, M. F. (2023). Removing the cardiac field artifact from the EEG using neural network regression. Psychophysiology, 60(10). 10.1111/PSYP.14323

Azevedo, R. T., Garfinkel, S. N., Critchley, H. D., & Tsakiris, M. (2017). ARTICLE Cardiac afferent activity modulates the expression of racial stereotypes. Nature Communications. 10.1038/ncomms13854

Azzalini, D., Rebollo, I., & Tallon-Baudry, C. (2019). Visceral Signals Shape Brain Dynamics and Cognition. In Trends in Cognitive Sciences (Vol. 23, Issue 6, pp. 488–509). Elsevier Ltd. 10.1016/j.tics.2019.03.007

Bai, Z., Fong, K. N. K., Zhang, J. J., Chan, J., & Ting, K. H. (2020). Immediate and long-term effects of BCI-based rehabilitation of the upper extremity after stroke: a systematic review and meta-analysis. Journal of Neuroengineering and Rehabilitation, 17(1). 10.1186/S12984-020-00686-2

Barrett, L. F., & Simmons, W. K. (2015). Interoceptive predictions in the brain. Nature Reviews Neuroscience 2015 16:7, 16(7), 419–429. 10.1038/nrn3950

Benjamini, Y., Krieger, A. M., & Yekutieli, D. (2006). Adaptive linear step-up procedures that control the false discovery rate. Biometrika, 93(3), 491–507. 10.1093/BIOMET/93.3.491

Bianchini, E., Mancuso, M., Zampogna, A., Guerra, A., & Suppa, A. (2021). Cardiac cycle does not affect motor evoked potential variability: A real-time EKG-EMG study. Brain Stimulation, 14(1), 170–172. 10.1016/J.BRS.2020.12.009

Biasiucci, A., Leeb, R., Iturrate, I., Perdikis, S., Al-Khodairy, A., Corbet, T., Schnider, A., Schmidlin, T., Zhang, H., Bassolino, M., Viceic, D., Vuadens, P., Guggisberg, A. G., & Millán, J. D. R. (2018). Brain-actuated functional electrical stimulation elicits lasting arm motor recovery after stroke. Nature Communications 2018 9:1, 9(1), 1–13. 10.1038/s41467-018-04673-z

Boudewyn, M. A., Luck, S. J., Farrens, J. L., & Kappenman, E. S. (2018). How Many Trials Does It Take to Get a Significant ERP Effect? It Depends. Psychophysiology, 55(6), e13049. 10.1111/PSYP.13049

Bury, G., García-Huéscar, M., Bhattacharya, J., & Ruiz, M. H. (2019). Cardiac afferent activity modulates early neural signature of error detection during skilled performance. NeuroImage, 199, 704–717. 10.1016/J.NEUROIMAGE.2019.04.043

Chettouf, S., Rueda-Delgado, L. M., de Vries, R., Ritter, P., & Daffertshofer, A. (2020). Are unimanual movements bilateral? Neuroscience & Biobehavioral Reviews, 113, 39–50. 10.1016/J.NEUBIOREV.2020.03.002

Cohen, M. X. (2019). A better way to define and describe Morlet wavelets for time-frequency analysis. NeuroImage, 199, 81–86. 10.1016/J.NEUROIMAGE.2019.05.048

Craig, A. D. (2009). How do you feel — now? The anterior insula and human awareness. Nature Reviews Neuroscience, 10(1), 59–70. 10.1038/nrn2555

Critchley, H. D., & Garfinkel, S. N. (2017). Interoception and emotion. Current Opinion in Psychology, 17, 7–14. 10.1016/J.COPSYC.2017.04.020

Critchley, H. D., & Garfinkel, S. N. (2018). The influence of physiological signals on cognition. Current Opinion in Behavioral Sciences, 19, 13–18. 10.1016/J.COBEHA.2017.08.014

Daly, A. E., & Bialocerkowski, A. E. (2009). Does evidence support physiotherapy management of adult Complex Regional Pain Syndrome Type One? A systematic review. *European Journal of Pain (London*, England), 13(4), 339–353. 10.1016/J.EJPAIN.2008.05.003

Destrieux, C., Fischl, B., Dale, A., & Halgren, E. (2010). Automatic parcellation of human cortical gyri and sulci using standard anatomical nomenclature. NeuroImage, 53(1), 1. 10.1016/J.NEUROIMAGE.2010.06.010

Duschek, S., Werner, N. S., & Reyes del Paso, G. A. (2013). The behavioral impact of baroreflex function: A review. Psychophysiology, 50(12), 1183–1193. 10.1111/PSYP.12136

Edwards, L., Ring, C., McIntyre, D., Carroll, D., & Martin, U. (2007). Psychomotor speed in hypertension: effects of reaction time components, stimulus modality, and phase of the cardiac cycle. Psychophysiology, 44(3), 459–468. 10.1111/J.1469-8986.2007.00521.X

Engelen, T., Solcà, M., & Tallon-Baudry, C. (2023). Interoceptive rhythms in the brain. Nature Neuroscience 2023, 1–15. 10.1038/s41593-023-01425-1

Filippi, M. M., Oliveri, M., Vernieri, F., Pasqualetti, P., & Rossini, P. M. (2000). Are autonomic signals influencing cortico-spinal motor excitability? A study with transcranial magnetic stimulation. Brain Research, 881(2), 159–164. 10.1016/S0006-8993(00)02837-7

Galvez-Pol, A., Virdee, P., Villacampa, J., & Kilner, J. M. (2022). Active tactile discrimination is coupled with and modulated by the cardiac cycle. ELife, 11. 10.7554/ELIFE.78126

Garfinkel, S. N., Minati, L., Gray, M. A., Seth, A. K., Dolan, R. J., & Critchley, H. D. (2014). Fear from the heart: sensitivity to fear stimuli depends on individual heartbeats. The Journal of Neuroscience : The Official Journal of the Society for Neuroscience, 34(19), 6573–6582. 10.1523/JNEUROSCI.3507-13.2014

Gray, M. A., Beacher, F. D., Minati, L., Nagai, Y., Kemp, A. H., Harrison, N. A., & Critchley, H. D. (2012). Emotional appraisal is influenced by cardiac afferent information. *Emotion (Washington*, D.C.), 12(1), 180–191. 10.1037/A0025083

Grissom, R. J., & Kim, J. J. (2012). Effect sizes for research: Univariate and multivariate applications, second edition. *Effect Sizes for Research: Univariate and Multivariate Applications*, Second Edition, 1–434. 10.4324/9780203803233

Grèzes, J., & Decety, J. (2001). Functional anatomy of execution, mental simulation, observation, and verb generation of actions: A meta-analysis. Human Brain Mapping, 12(1), 1. 10.1002/1097-0193(200101)12:1<1::aid-hbm10>3.0.co;2-v

Grund, M., Al, E., Pabst, M., Dabbagh, A., Stephani, T., Nierhaus, T., Gaebler, M., & Villringer, A. (2022). Respiration, Heartbeat, and Conscious Tactile Perception. Journal of Neuroscience, 42(4), 643–656. 10.1523/JNEUROSCI.0592-21.2021

Hardwick, R. M., Caspers, S., Eickhoff, S. B., & Swinnen, S. P. (2018). Neural correlates of action: Comparing meta-analyses of imagery, observation, and execution. In Neuroscience and Biobehavioral Reviews (Vol. 94, pp. 31–44). Elsevier Ltd. 10.1016/j.neubiorev.2018.08.003

Helin, P., Sihvonen, T., & Hänninen, O. (1987). Timing of the triggering action of shooting in relation to the cardiac cycle. British Journal of Sports Medicine, 21(1), 33–36. 10.1136/BJSM.21.1.33

Hétu, S., Grégoire, M., Saimpont, A., Coll, M. P., Eugène, F., Michon, P. E., & Jackson, P. L. (2013). The neural network of motor imagery: An ALE meta-analysis. Neuroscience & Biobehavioral Reviews, 37(5), 930–949. 10.1016/J.NEUBIOREV.2013.03.017

Honda, T., & Nakao, T. (2022). Impact of Cardiac Interoception on the Self-Prioritization Effect. Frontiers in Psychology, 13, 825370. 10.3389/FPSYG.2022.825370/BIBTEX

Hyvärinen, A., & Oja, E. (2000). Independent component analysis: algorithms and applications. Neural Networks : The Official Journal of the International Neural Network Society, 13(4–5), 411–430. 10.1016/S0893-6080(00)00026-5

Khalsa, S. S., Adolphs, R., Cameron, O. G., Critchley, H. D., Davenport, P. W., Feinstein, J. S., Feusner, J. D., Garfinkel, S. N., Lane, R. D., Mehling, W. E., Meuret, A. E., Nemeroff, C. B., Oppenheimer, S., Petzschner, F. H., Pollatos, O., Rhudy, J. L., Schramm, L. P., Simmons, W. K., Stein, M. B., … Zucker, N. (2018). Interoception and Mental Health: A Roadmap. Biological Psychiatry. Cognitive Neuroscience and Neuroimaging, 3(6), 501–513. 10.1016/J.BPSC.2017.12.004

Khan, M. A., Das, R., Iversen, H. K., & Puthusserypady, S. (2020). Review on motor imagery based BCI systems for upper limb post-stroke neurorehabilitation: From designing to application. In Computers in Biology and Medicine (Vol. 123, p. 103843). Elsevier Ltd. 10.1016/j.compbiomed.2020.103843

Khoury, N. M., Lutz, J., & Schuman-Olivier, Z. (2018). Interoception in Psychiatric Disorders: A Review of Randomized Controlled Trials with Interoception-based Interventions. Harvard Review of Psychiatry, 26(5), 250. 10.1097/HRP.0000000000000170

Klein, A., & Tourville, J. (2012). 101 labeled brain images and a consistent human cortical labeling protocol. Frontiers in Neuroscience, 6(DEC). 10.3389/FNINS.2012.00171

Konrad, P. (2005). The ABC of EMG.

Konttinen, N., Mets, T., Lyytinen, H., & Paananen, M. (2003). Timing of triggering in relation to the cardiac cycle in nonelite rifle shooters. Research Quarterly for Exercise and Sport, 74(4), 395–400. 10.1080/02701367.2003.10609110

Kunzendorf, S., Klotzsche, F., Akbal, M., Villringer, A., Ohl, S., & Gaebler, M. (2019). Active information sampling varies across the cardiac cycle. Psychophysiology, 56(5), e13322. 10.1111/PSYP.13322

Kurbel, S. (2014). A vector-free ECG interpretation with P, QRS & T waves as unbalanced transitions between stable configurations of the heart electric field during P-R, S-T & T-P segments. Theoretical Biology and Medical Modelling, 11(1), 1–21.

Lacey, J. (1967). Somatic response patterning and stress : some revisions of activation theory.

Li, H., Huang, G., Lin, Q., Zhao, J. L., Lo, W. L. A., Mao, Y. R., Chen, L., Zhang, Z. G., Huang, D. F., & Li, L. (2018). Combining Movement-Related Cortical Potentials and Event-Related Desynchronization to Study Movement Preparation and Execution. Frontiers in Neurology, 9(OCT), 822. 10.3389/FNEUR.2018.00822

Lotze, M., Montoya, P., Erb, M., Hülsmann, E., Flor, H., Klose, U., Birbaumer, N., & Grodd, W. (1999). Activation of Cortical and Cerebellar Motor Areas during Executed and Imagined Hand Movements: An fMRI Study. http://direct.mit.edu/jocn/article-pdf/11/5/491/1758586/089892999563553.pdf

Makowski, D., Pham, T., Lau, Z. J., Brammer, J. C., Lespinasse, F., Pham, H., Schölzel, C., & Chen, S. H. A. (2021). NeuroKit2: A Python toolbox for neurophysiological signal processing. Behavior Research Methods, 53(4), 1689–1696. 10.3758/S13428-020-01516-Y/TABLES/3

Malouin, F., Richards, C. L., Jackson, P. L., Lafleur, M. F., Durand, A., & Doyon, J. (2007). The kinesthetic and visual imagery questionnaire (KVIQ) for assessing motor imagery in persons with physical disabilities: A reliability and construct validity study. Journal of Neurologic Physical Therapy, 31(1), 20–29. 10.1097/01.NPT.0000260567.24122.64

Marozzi, M. (2004). Some remarks about the number of permutations one should consider to perform a permutation test. Statistica, 64(1), 193–201. 10.6092/ISSN.1973-2201/32

Motyka, P., Grund, M., Forschack, N., Al, E., Villringer, A., & Gaebler, M. (2019). Interactions between cardiac activity and conscious somatosensory perception. Psychophysiology, 56(10), e13424. 10.1111/PSYP.13424

Neuper, C., Scherer, R., Reiner, M., & Pfurtscheller, G. (2005). Imagery of motor actions: differential effects of kinesthetic and visual-motor mode of imagery in single-trial EEG. Brain Research. Cognitive Brain Research, 25(3), 668–677. 10.1016/J.COGBRAINRES.2005.08.014

Nord, C. L., & Garfinkel, S. N. (2022). Interoceptive pathways to understand and treat mental health conditions. Trends in Cognitive Sciences, 26(6), 499–513. 10.1016/J.TICS.2022.03.004

Ohl, S., Wohltat, C., Kliegl, R., Pollatos, O., & Engbert, R. (2016). Microsaccades Are Coupled to Heartbeat. The Journal of Neuroscience : The Official Journal of the Society for Neuroscience, 36(4), 1237–1241. 10.1523/JNEUROSCI.2211-15.2016

Otsuru, N., Miyaguchi, S., Kojima, S., Yamashiro, K., Sato, D., Yokota, H., Saito, K., Inukai, Y., & Onishi, H. (2020). Timing of Modulation of Corticospinal Excitability by Heartbeat Differs with Interoceptive Accuracy. Neuroscience, 433, 156–162. 10.1016/j.neuroscience.2020.03.014

Park, H. D., Barnoud, C., Trang, H., Kannape, O. A., Schaller, K., & Blanke, O. (2020). Breathing is coupled with voluntary action and the cortical readiness potential. Nature Communications, 11(1). 10.1038/S41467-019-13967-9

Park, H. D., & Tallon-Baudry, C. (2014). The neural subjective frame: From bodily signals to perceptual consciousness. Philosophical Transactions of the Royal Society B: Biological Sciences, 369(1641). 10.1098/rstb.2013.0208

Park, H.-D., Correia, S., Ducorps, A., & Tallon-Baudry, C. (2014). Spontaneous fluctuations in neural responses to heartbeats predict visual detection. 17(4). 10.1038/nn.3671

Peirce, J., Gray, J. R., Simpson, S., MacAskill, M., Höchenberger, R., Sogo, H., Kastman, E., & Lindeløv, J. K. (2019). PsychoPy2: Experiments in behavior made easy. Behavior Research Methods, 51(1), 195–203. 10.3758/S13428-018-01193-Y/FIGURES/3

Perrin, F., Pernier, J., Bertrand, O., & Echallier, J. F. (1989). Spherical splines for scalp potential and current density mapping. Electroencephalography and Clinical Neurophysiology, 72(2), 184–187. 10.1016/0013-4694(89)90180-6

Pichiorri, F., Morone, G., Petti, M., Toppi, J., Pisotta, I., Molinari, M., Paolucci, S., Inghilleri, M., Astolfi, L., Cincotti, F., & Mattia, D. (2015). Brain–computer interface boosts motor imagery practice during stroke recovery. Annals of Neurology, 77(5), 851–865. 10.1002/ANA.24390

Porro, C. A., Cettolo, V., Francescato, M. P., & Baraldi, P. (2000). Ipsilateral involvement of primary motor cortex during motor imagery. European Journal of Neuroscience, 12(8), 3059–3063. 10.1046/J.1460-9568.2000.00182.X

Pramme, L., Larra, M. F., Schächinger, H., & Frings, C. (2014). Cardiac cycle time effects on mask inhibition. Biological Psychology, 100(1), 115–121. 10.1016/J.BIOPSYCHO.2014.05.008

Pramme, L., Larra, M. F., Schächinger, H., & Frings, C. (2016). Cardiac cycle time effects on selection efficiency in vision. Psychophysiology, 53(11), 1702–1711. 10.1111/PSYP.12728

Rae, C. L., Botan, V. E., Gould Van Praag, C. D., Herman, A. M., Nyyssönen, J. A. K., Watson, D. R., Duka, T., Garfinkel, S. N., & Critchley, H. D. (2018). Response inhibition on the stop signal task improves during cardiac contraction. Scientific Reports 2018 8:1, 8(1), 1–9. 10.1038/s41598-018-27513-y

Ramos-Murguialday, A., Broetz, D., Rea, M., Läer, L., Yilmaz, Ö., Brasil, F. L., Liberati, G., Curado, M. R., Garcia-Cossio, E., Vyziotis, A., Cho, W., Agostini, M., Soares, E., Soekadar, S., Caria, A., Cohen, L. G., & Birbaumer, N. (2013). Brain-machine interface in chronic stroke rehabilitation: a controlled study. Annals of Neurology, 74(1), 100–108. 10.1002/ANA.23879

Rau, H., Pauli, P., Brody, S., Elbert, T., & Birbaumer, N. (1993). Baroreceptor stimulation alters cortical activity. Psychophysiology, 30(3), 322–325. 10.1111/J.1469-8986.1993.TB03359.X

Ruscio, J., & Mullen, T. (2012). Confidence Intervals for the Probability of Superiority Effect Size Measure and the Area Under a Receiver Operating Characteristic Curve. Multivariate Behavioral Research, 47(2), 201–223. 10.1080/00273171.2012.658329

Rimbert, S., & Fleck, S. (2023). Long-term kinesthetic motor imagery practice with a BCI: Impacts on user experience, motor cortex oscillations and BCI performances. Computers in Human Behavior, 146, 107789. 10.1016/J.CHB.2023.107789

Sannelli, C., Vidaurre, C., Müller, K.-R., & Blankertz, B. (2019). A large scale screening study with a SMR-based BCI: Categorization of BCI users and differences in their SMR activity. PLOS ONE, 14(1), e0207351. 10.1371/JOURNAL.PONE.0207351

Savaki, H. E., & Raos, V. (2019). Action perception and motor imagery: Mental practice of action. Progress in Neurobiology, 175, 107–125. 10.1016/J.PNEUROBIO.2019.01.007

Schulz, A., Reichert, C. F., Richter, S., Lass-Hennemann, J., Blumenthal, T. D., & Schächinger, H. (2009). Cardiac modulation of startle: Effects on eye blink and higher cognitive processing. Brain and Cognition, 71(3), 265–271. 10.1016/J.BANDC.2009.08.002

Sekihara, K., & Nagarajan, S. S. (2008). Adaptive Spatial Filters for Electromagnetic Brain Imaging. Adaptive Spatial Filters for Electromagnetic Brain Imaging. 10.1007/978-3-540-79370-0

Tallon-Baudry, C., Bertrand, O., Delpuech, C., & Pernier, J. (1997). Oscillatory-Band (30-70 Hz) Activity Induced by a Visual Search Task in Humans.

Tong, Y., Pendy, J. T., Li, W. A., Du, H., Zhang, T., Geng, X., & Ding, Y. (2016). Motor Imagery-Based Rehabilitation: Potential Neural Correlates and Clinical Application for Functional Recovery of Motor Deficits after Stroke. 10.14336/AD.2016.1012

Van Veen, B. D., Van Drongelen, W., Yuchtman, M., & Suzuki, A. (1997). Localization of brain electrical activity via linearly constrained minimum variance spatial filtering. IEEE Transactions on Biomedical Engineering, 44(9), 867–880. 10.1109/10.623056

van Vliet, M., Liljeström, M., Aro, S., Salmelin, R., & Kujala, J. (2018). Analysis of functional connectivity and oscillatory power using DICS: From Raw MEG data to group-level statistics in python. Frontiers in Neuroscience, 12(SEP). 10.3389/fnins.2018.00586

Yang, Y. J., Jeon, E. J., Kim, J. S., & Chung, C. K. (2021). Characterization of kinesthetic motor imagery compared with visual motor imageries. Scientific Reports 2021 11:1, 11(1), 1–11. 10.1038/s41598-021-82241-0

