## Supplementary material for "Cardiac Cycle Modulates Alpha and Beta Suppression during Motor Imagery": Figure S1, Figure S2, Figure S3, Figure S4, Table S1, Table S2, Table S3, Table S4

### EMG Activity

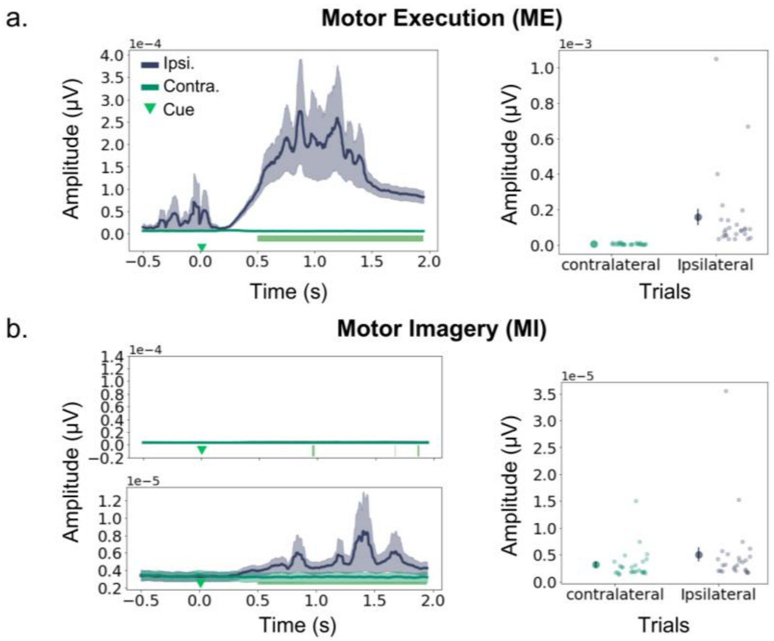


**Figure S1. Differences between Ipsilateral and Contralateral EMG waveforms.** Same as **Figure 3ab** but including all n=26 participants for which EMG was available. Statistical differences are as reported in the main text.

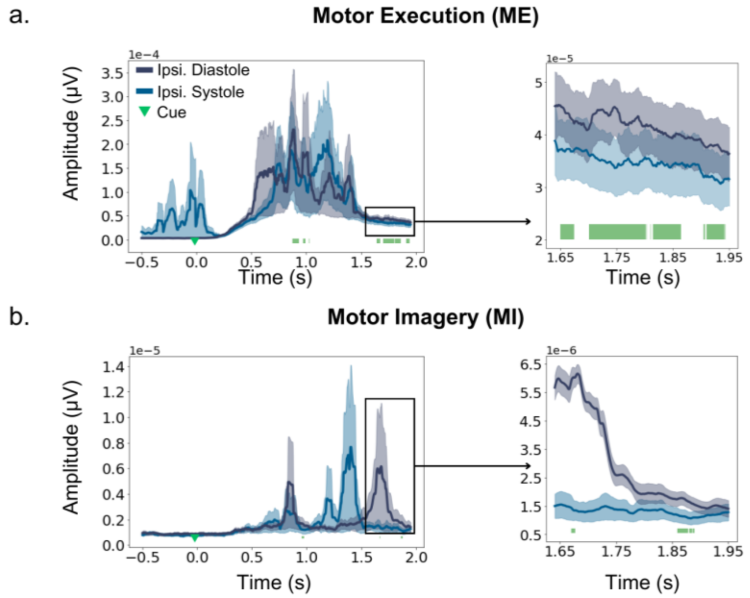

**Figure S2. Differences between Ipsilateral EMG waveforms for Systole-cued and Diastole-cued Trials.** Same as panels in **Figure 3cd** but including all n=26 participants for which EMG was available. Statistical differences are as reported in the main text and caption to Figure 3cd.

#### Source Analysis: Full-epoch Length

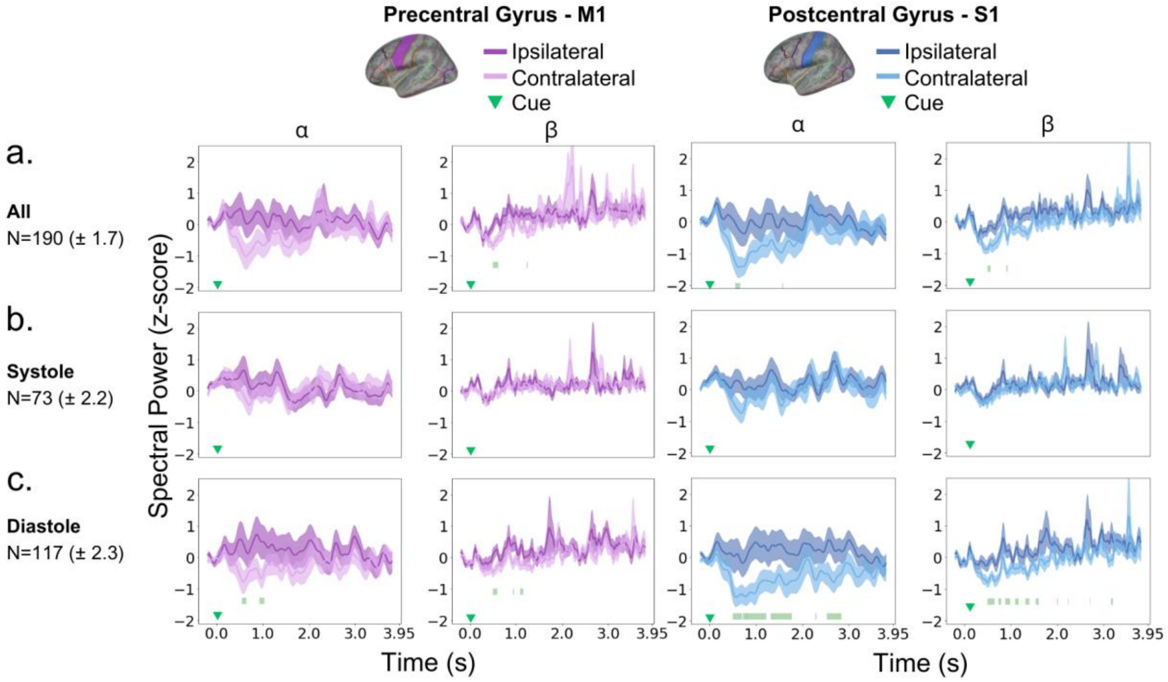


**Figure S3. Differences between Ipsilateral and Contralateral Power Spectral Density within the full-epoch length.** This figure shows the results of our exploratory analysis on source-reconstructed epochs from 0.5 to 4 seconds after the cue. The panels are similar to panels in Figure 4 but extending up to 4 seconds post-cue.

#### Cardiac Alignment

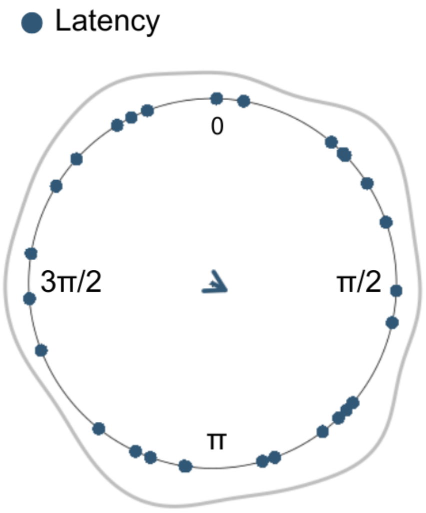


**Figure S4. Assessing the Cardiac to Cue Alignment Hypothesis.** The figure shows the results of the Rayleigh test to assess whether participants aligned their heartbeat with the presentation of the cue. As shown by the resultant mean vector (blue arrow), there was no deviation from the uniform distribution (Z=0.0773, P=0.8483).

#### Control Analysis on Source-reconstructed Epochs

**Table S1. Control analysis in S1 for all trials.** Contralateral versus ipsilateral suppression in alpha and beta bands in S1 and M1 was evaluated across subsets of trials (x10 times: 10 control runs) from the total set of trials, independent of cardiac phases, with the number of trials matched to the systole subset. The *p-value* ranges for statistical differences at each time point within the assessed window, 0.5–4 s, are provided. Significant effects after FDR control are marked with an asterisk *, and are accompanied by non-parametric effect sizes, $\Delta_{dep}$.

| **Control runs** | **All Trials** | | | |
|---|---|---|---|---|
| | **Alpha** | | **Beta** | |
| | **P-value ($P_{FDR}$)** | **Effect size ($\Delta_{dep}$)** | **P-value ($P_{FDR}$)** | **Effect size ($\Delta_{dep}$)** |
| **1** | 0.001–0.9926 | / | 0.0001–0.0001* | 0.79 |
| **2** | 0.0006–0.0094* | 0.76 | 0.0001–0.0072* | 0.83 |
| **3** | 0.0268–0.0997 | / | 0.0012–0.9778 | / |
| **4** | 0.0001–0.0098* | 0.86 | 0.0026–0.0102* | 0.76 |
| **5** | 0.0002–0.0058* | 0.86 | 0.0086–0.9742 | / |
| **6** | 0.0002–0.0008* | 0.90 | 0.002–0.9988 | / |
| **7** | 0.0006–0.006* | 0.76 | 0.0001–0.0038* | 0.76 |
| **8** | 0.0532–0.951 | / | 0.001–0.991 | / |
| **9** | 0.0276–0.9862 | / | 0.0008–0.9966 | / |
| **10** | 0.0022–0.9946 | / | 0.0028–0.993 | / |

*Note: * Significant after FDR*

**Table S2. Control analysis in S1 for all diastole-cued trials.** Same as Table S1 but for subsets of diastole-cued trials, assessing the consistency of significant contralateral suppression when cues instructing movement direction occur in the diastolic phase of the cardiac cycle.

| **Control runs** | **Diastole** | | | |
|---|---|---|---|---|
| | **Alpha** | | **Beta** | |
| | **P-value ($P_{FDR}$)** | **Effect size ($\Delta_{dep}$)** | **P-value ($P_{FDR}$)** | **Effect size ($\Delta_{dep}$)** |
| **1** | 0.0001–0.0372* | 0.86 | 0.0008–0.0438* | 0.86 |
| **2** | 0.0001–0.0254* | 0.90 | 0.0001–0.0256* | 0.83 |
| **3** | 0.0014–0.9986 | / | 0.001–0.9584 | / |
| **4** | 0.0001–0.0552* | 0.83 | 0.0016–0.0524* | 0.76 |
| **5** | 0.0001–0.0268* | 0.76 | 0.0004–0.0292* | 0.90 |
| **6** | 0.0004–0.0176* | 0.80 | 0.0008–0.0196* | 0.83 |
| **7** | 0.0001–0.0296* | 0.83 | 0.0001–0.0336* | 0.90 |
| **8** | 0.0001–0.0124* | 0.79 | 0.0004–0.0132* | 0.79 |
| **9** | 0.0001–0.0262* | 0.90 | 0.0002–0.027* | 0.83 |
| **10** | 0.0001–0.0342* | 0.83 | 0.0008–0.0344* | 0.83 |

*Note: * Significant after FDR*

**Table S3. Control analysis in M1 for all trials.** Same as Table S1 but in M1.

| **Control runs** | **All Trials** | | | |
|---|---|---|---|---|
| | **Alpha** | | **Beta** | |
| | **P-value ($P_{FDR}$)** | **Effect size ($\Delta_{dep}$)** | **P-value ($P_{FDR}$)** | **Effect size ($\Delta_{dep}$)** |
| **1** | 0.0022–0.9992 | / | 0.0004–0.9944 | / |
| **2** | 0.0032–0.0072* | 0.76 | 0.0002–0.0092* | 0.83 |
| **3** | 0.0014–1 | / | 0.0014–0.9998 | / |
| **4** | 0.0002–0.0104* | 0.83 | 0.0004–0.0088* | 0.72 |
| **5** | 0.0354–0.9508 | / | 0.0226–0.9972 | / |
| **6** | 0.0076–0.9742 | / | 0.0068–0.9778 | / |
| **7** | 0.0496–0.9946 | / | 0.0001–0.006* | 0.79 |
| **8** | 0.037–0.9422 | / | 0.0001–0.9858 | / |
| **9** | 0.1192–0.9852 | / | 0.006–0.9998 | / |
| **10** | 0.0118–0.9914 | / | 0.0064–0.9948 | / |

*Note: * Significant after FDR*

**Table S4. Control analysis in M1 for diastole-cued trials.** Same as Table S1 but for M1 and diastole-cued trials.

| Control runs | Diastole | | | |
|---|---|---|---|---|
| | Alpha | | Beta | |
| | P-value ($P_{FDR}$) | Effect size ($\Delta_{dep}$) | P-value ($P_{FDR}$) | Effect size ($\Delta_{dep}$) |
| **1** | 0.0018–0.0424* | 0.83 | 0.012–0.043* | 0.79 |
| **2** | 0.0038–0.0256* | 0.83 | 0.0158–0.0242* | 0.83 |
| **3** | 0.0036–0.9966 | / | 0.0192–0.9924 | / |
| **4** | 0.0001–0.0476* | 0.86 | 0.0084–0.0508* | 0.72 |
| **5** | 0.004–0.0306* | 0.76 | 0.0001–0.0304* | 0.90 |
| **6** | 0.0006–0.02* | 0.83 | 0.0114–0.0194* | 0.59 |
| **7** | 0.0006–0.0336* | 0.79 | 0.0016–0.032* | 0.79 |
| **8** | 0.0012–0.0116* | 0.72 | 0.0356–0.98 | / |
| **9** | 0.001–0.0266* | 0.83 | 0.0158–0.0274* | 0.69 |
| **10** | 0.0076–0.0346* | 0.76 | 0.0018–0.0338* | 0.79 |

*Note: * Significant after FDR*
